# Idiosyncratic and generic single nuclei and spatial transcriptional patterns in papillary and anaplastic thyroid cancers

**DOI:** 10.1101/2024.02.15.580495

**Authors:** Adrien Tourneur, Joel Rodrigues Vitória, Manuel Saiselet, Ligia Craciun, Denis Larsimont, Anne Lefort, Frederick Libert, Carine Maenhaut, Sabine Costagliola, Maxime Tarabichi, Mirian Romitti, Vincent Detours

**Affiliations:** IRIBHM Jacques E. Dumont, Université Libre de Bruxelles (ULB), 808 route de Lennik, 1070 Brussels, Belgium; Department of Pathology, Institut Jules Bordet, Rue Meylemeersch 90, 1070 Bruxelles, Belgium; Interuniversity Institute of Bioinformatics in Brussels ULB-VUB (IB2), La Plaine Campus, Blv du Triomphe, 1050 Brussels, Belgium

## Abstract

Sixty percent of papillary thyroid cancers (PTCs) are driven by *BRAF^V600E^*, a mutation associated with high inter- and intra-tumoral heterogeneity. PTCs may become highly aggressive anaplastic thyroid cancers (ATC). While single cell transcriptomics may resolve this heterogeneity, it is potentially confounded by technical effects whose correction may dampen inter-tumor variations. Here we profiled ATCs and *BRAF^V600E^* PTCs with single nuclei RNA-seq and spatial transcriptomics, and an experimental design disentangling biological and technical variations. It reveals that much transcriptional variation in cancer cells and several immune cell types is idiosyncratic, i.e. tumor-specific, a phenomenon obscured by batch integration in a number of single cell studies. Idiosyncrasies are associated in some cases with genomic aberrations and global tissue states like hypoxia. Beyond idiosyncrasies, differentiation markers *SLC5A5* (*NIS*), *TPO*, *TG* and *TSHR* are lost in a sequence mirrored by their gain during human thyroid organoids maturation, suggesting a new classification of cancer cell states. PTC cells retain *TSHR* expression and show features of partial EMT with a massive expression of *FN1*, which promotes proliferation via an autocrine loop. In contrast, ATCs undergo full blown EMT, with expression of mesenchymal extracellular components and loss of *TSHR*. Finally, we show that the microenvironment of cancer cells is driven by inflammation. These findings may help future stratifications of *BRAF^V600E^* PTCs.

## Introduction

Thyroid cancer (TC) is the most common endocrine malignancy (1) and its incidence is expected to keep rising (2). Papillary thyroid cancer (PTC) is the most common form of TC, accounting for 70% to 90% of cases (3–6). Anaplastic thyroid cancer (ATC) is a very rare form of TC but is considered among the most lethal human cancers (4, 7). It is often associated with low tumor purity, and especially high macrophage infiltration (8). 60% of PTC bear the *BRAF^V600E^* mutation which is associated with some clinical aggressiveness features. *BRAF^V600E^* mutated PTCs present variations in their clinical features and tumor purity, with a significant fraction of them having low tumor purity and a highly cellular tumor microenvironment (9). It is thus challenging to stratify this disease and find the optimal treatment option for patients, and a better classification would bring clinical benefits (10).

The Cancer Genome Atlas (TCGA) consortium profiled 496 PTC samples using different DNA and RNA sequencing technologies, which all relied on bulk sequencing (11). This large cohort provided a landscape of the inter-patient heterogeneity in PTC. One new concept it brought forth is the gene-expression-based *BRAF^V600E^*-*RAS* score (BRS). It provides a way to position tumors on a quantitative scale associated with some clinical features. It also varies within tumors bearing the *BRAF^V600E^* mutation, once again highlighting the heterogeneity of this disease. Yet, bulk methods are not suited to fully understand and stratify low purity tumors or subclonally diverse ones. Single cell RNA sequencing (scRNA-seq) can tackle low purity and, to some extent, subclonal diversity, in tumor samples.

But the complex data it generates requires sophisticated data analysis which dramatically impacts subsequent cell stratification. In particular, in most scRNA-seq studies, each sample is processed in an individual experiment, with its specific, unknown, batch effects. These are typically removed with batch effect correction algorithms. Those methods operate a delicate balance between biological signal preservation and technical noise removal that relies on some assumptions that must be verified (12, 13). But current experimental designs do not disentangle technical and biological inter-sample variation. Thus, there is a real possibility that batch correction also dampens genuine biological differences between samples. Out of the 12 publications we found with scRNA-seq data for TC, one includes only ATC samples (14), two include both ATC and PTC (7, 15), and 10 include only PTC (5, 6, 16–24), along with other tissue types for some (e.g. adjacent normal, metastatic, etc.). Four of those 12 publications mention using batch effect correction algorithms, but discuss very little the consequences they have on biological signal and cell stratification (7, 17, 20, 24). Here we propose an experimental design that disentangles batch effects from biological variations. We find that technical variation is negligible in our data, and that transcriptional states of several epithelial and non epithelial cell types are idiosyncratic, i.e. tumor-specific.

Some of the above publications address the classification of epithelial cells in various contexts. scRNA-seq studies generally include low amounts of samples, which can hinder classification attempts of disease categories that are too broad. To narrow down the disease categories we address while still studying respectively the most prevalent and aggressive forms of the disease, we decided to address this classification in the context of *BRAF^V600E^* PTC and to ATC samples. We studied 16 samples of primary thyroid cancers ranging 4 different subtypes. Previous thyroid cancer studies relying on scRNA-seq presented cohorts of 5 to 22 thyroid cancer samples ranging 1 to 4 different subtypes.

Combining the respective strengths of multiplex immunofluorescence (MIF), single nuclei RNA sequencing (snRNA-seq), spatial RNA sequencing (spRNA-seq) and the TCGA, we provide new insights into ATC and PTC heterogeneity at whole transcriptome and cellular resolution with spatial context, establishing links between transcriptomics and the H&E images routinely used in the clinics. This dataset spans the spectrum of thyroid dedifferentiation, from fully differentiated hormone-producing thyrocytes to fully dedifferentiated mesenchymal-like ATC cancer cells. We propose a new classification for cells of epithelial origin in thyroid cancer.

## Results

### Sample-match single nuclei and spatial transcriptomes of papillary and anaplastic thyroid cancers

We profiled 10 PTC samples (including classical, warthin-like and follicular variants) from 10 patients harboring the *BRAF^V600E^* mutation and 6 ATC samples from 4 patients with mutations in *TP53*, *PTEN*, *RB1* and *HRAS* (Supplemental Table 1). The *BRAF^V600E^* mutation has been shown to be clonal in PTC (9, 11).

Our profiling protocol integrated adjacent spRNA-seq, snRNA-seq and MIF in the same frozen blocks. Each 10 μm-thick spRNA-seq slice was sandwiched by three 6 μm MIF slices on both sides, and those series of sandwiches were surrounded by two series of 20 μm snRNA-seq slices. MIF and spRNA-seq slices were cut consecutively to be immediately adjacent and to obtain a comparable tissue composition between adjacent slices, allowing them to be compared. Tumors in this cohort were resected in separate patients, except for two blocks, ATC3A-ATC3B and ATC4A-ATC4B, that were located ∼1cm apart in the same tumors. These four samples were used for four different snRNA-seq experiments and two different spRNA-seq experiments (ATC4A-B were not profiled with spRNA-seq).

Overall, the snRNA-seq dataset included 91,632 nuclei with a median of 1,415 genes and 2,337 UMI per nuclei (Supplemental Table 2). The spRNA-seq dataset included seven slides containing each three to six tissue slices processed in parallel as separate samples. In total 37 tissue slices were profiled, with 25 slices coming from PTC and 12 from ATC samples. These 37 slices contained on average 514 spots covered by the tissue. The median UMI and gene counts per spot were 7,670 and 2,900, respectively (Supplemental Table 3).

### Sample mixing experiments demonstrate idiosyncratic epithelial and non-epithelial cell states

Applying a standard normalization (SCTransform (25)) on our initial snRNA-seq dataset revealed that 9 clusters out of the 32 contained more than 90% nuclei from one particular sample (Figure 1A). As in most single cell studies, a single sample was processed in each experiment, raising a question: are these tumor-specific clusters the result of genuine biological variation between tumors or the result of tumor-specific experimental batch effects?

**Figure 1.**
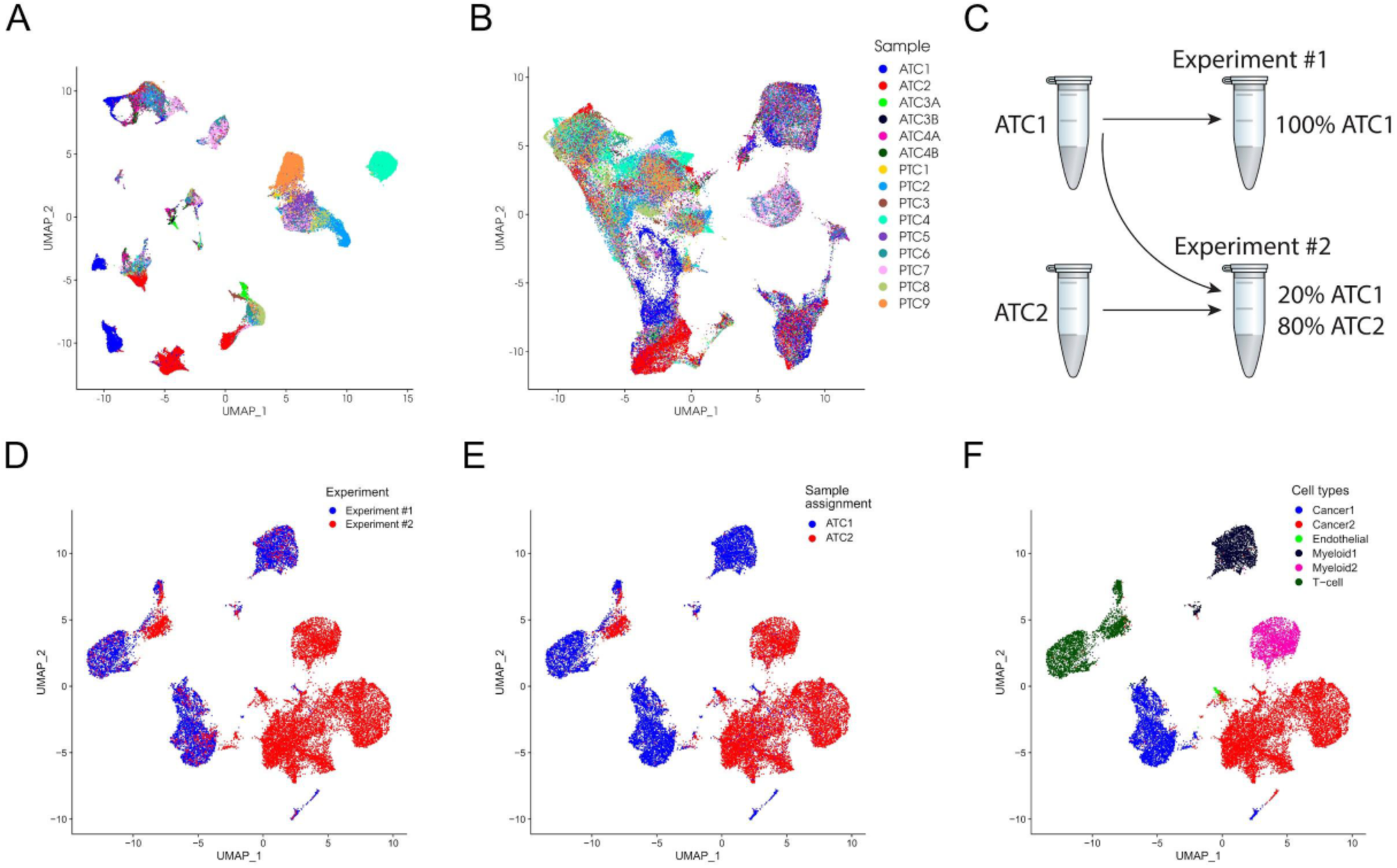
Idiosyncratic single nuclei profiles. **(A)** UMAP of nuclei transcriptomes from 9 PTC and 6 ATC samples without batch integration. **(B)** UMAP of the dataset after harmony batch integration. Integration drastically reduces the number of sample-specific clusters. **(C)** Schematic representation of the nuclei mixing experiment for samples ATC1 and ATC2. **(D)** UMAP showing the experimental tube of origin for experiments ATC1SN1 and ATC2SN1. Nuclei from Experiment #1, but not those from Experiment #2 are present in all clusters. **(E)** UMAP showing the sample of origin of each nuclei as assigned by Vireo. All clusters are tumor-specific. Thus, nuclei cluster by tumor, not by experiment. **(F)** UMAP showing the cell type annotation for samples ATC1 and ATC2.

Many studies have assumed the latter and applied computational batch effect removal (e.g. (7, 17, 24)), also known as data integration. To explore the effect of this approach we ran three popular data integration algorithms, including Seurat’s canonical correlation analysis and anchor-based merging (26), and Harmony (27). They all massively dampened inter-sample differences (Figure 1B, Supplemental Figure 1A-D) with all samples homogeneously distributed in every cluster. This is the desired effect if most of the variation is technical. If, on the contrary, most of the variation is biological, data integration would remove it, leading to misleading conclusions, for example that cells from different PTCs form a transcriptionally homogenous group.

We performed a sample mixing experiment to disentangle the contributions of technical and biological variations to sample-specific clusters. SNPs unique to individual patients were extracted from their ATC1 and ATC2 spRNA-seq data. These SNPs were then used to recover patient/tumor identity with Vireo (8) from a scRNA-seq experiment in which we mixed cells from ATC1 and ATC2 (Experiment #2 in Figure 1C). Another experiment included exclusively cells from ATC1 (Experiment #1 in Figure 1C).

Cells from the two experiments were merged as one dataset and clustered—without prior batch effect removal. Only 2% of cells from Experiment #1 were assigned to ATC2 by Vireo, establishing the accuracy of cells’ patient/tumor assignment. Coloring cells by experiment revealed that a set of cluster contained exclusively cells from Experiment #1, while clusters containing cells from Experiment #2 also included cells from Experiment #1 (Figure 1D). By contrast, coloring cells by patient/tumor revealed that all clusters were almost patient/tumor-specific (Figure 1E). Thus, clusters were associated with patient/tumor, not with experimental variation. The minimal batch effect is consistent with our parallel handling of cells profiled in this study in the same 10X Chromium run.

Cancer cells formed patient-specific clusters (Figures 1E-F). More surprisingly, myeloid and to a lesser extent T cells also grouped in patient-specific clusters.

Overall, this patient mixing experiment demonstrated that batch effects had a minimal contribution to transcriptome variation compared to biological effects. Transcriptional patient/tumor’s idiosyncratic states were present in both cancer and non cancer cells. As the core hypothesis underlying batch effect removal was clearly violated, we did not apply it in subsequent analyses.

### The micro-environnement is idiosyncratic in size and spatial distribution and is associated by systemic features such as thyroiditis or hypoxia

To capture the branching and hierarchical nature of differentiation, we annotated cell types at low, medium and high (sub)clustering granularities (details in Material and Methods), yielding partitions of cells into 11, 16 and 53 cell types, respectively (Figure 2A, Supplemental Figures 2A and 2C, cell types markers in Supplemental Tables 4-6). The medium granularity annotation, Figure 2A, is used in this study unless specified otherwise.

**Figure 2.**
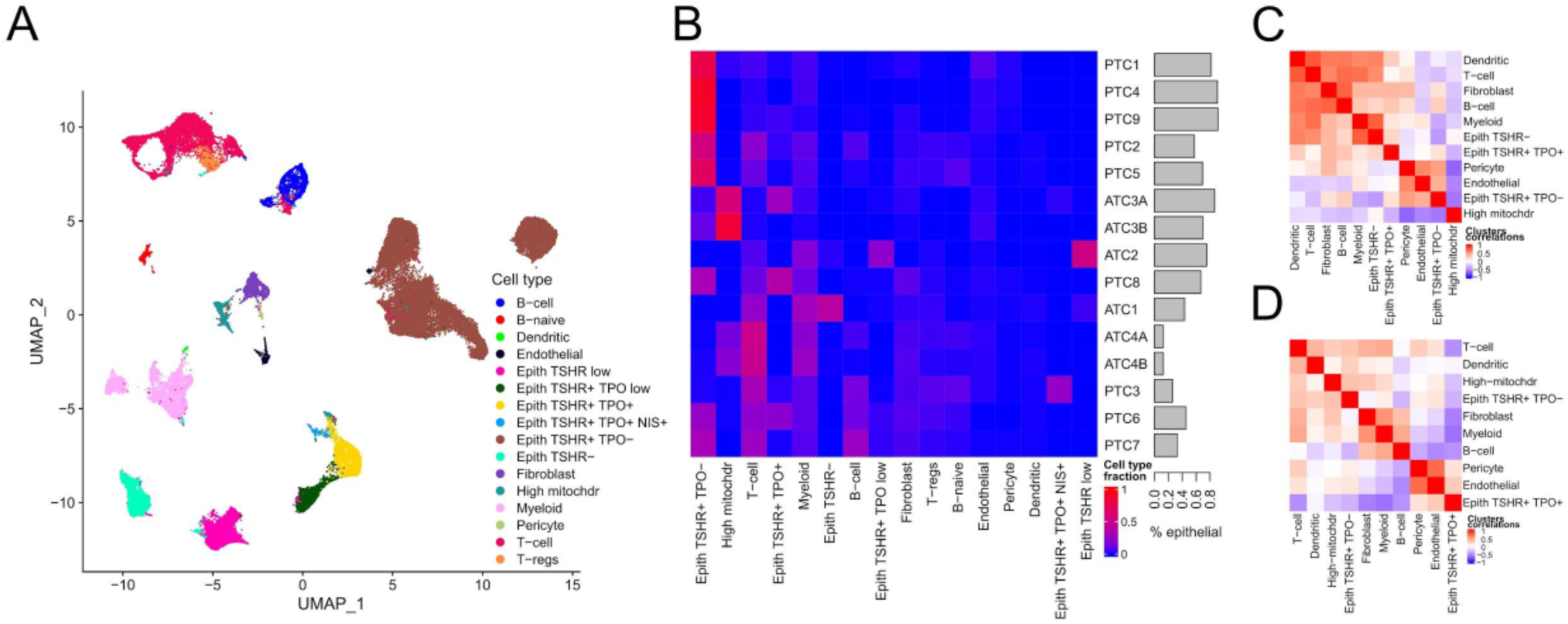
Idiosyncratic microenvironments. **(A)** UMAP of 9 PTC and 6 ATC samples with cell type annotation at medium resolution. **(B)** Cell type proportions per sample (left panel) and barplot of percentages of nuclei of epithelial origin in snRNA-seq dataset (right panel). **(C)** Correlogram of cell type clusters proportions in snRNA-seq dataset. **(D)** Correlogram of cell type clusters in 234 *BRAF^V600E^* PTCs from TCGA. Row and column in panel B-D were ordered with hierarchical clustering.

The size of the microenvironment showed remarkable variations in our dataset, ranging from 8.4 to 87.5% of the cells (Figure 2B). A clustering of sample-wise cell type proportions revealed three categories of tumors: 1- ATC that have the most idiosyncratic compositions, 2- PTC mainly composed of nuclei of epithelial origin and 3- PTC with a large immune compartment. To visualize the co-occurrence of different cell types we computed correlations between cell types proportions across samples in our dataset (Figure 2C), and as a validation, in 234 *BRAF^V600E^* PTCs from TCGA (Figure 2D, details in Material and Methods). Three groups of coordinated cell types were clearly visible. The first reflects the known association of inflammatory and mesenchymal cells in desmoplasia (9) : immune nuclei and fibroblasts co-occurred in the same samples, (Spearman correlation ρ = 0.77 for fibroblasts and B cells and 0.61 for fibroblasts and T cells in our dataset, in the TCGA cohort 0.42 and 0.43 respectively). The second cell type group reflected the association of EMT nuclei, which are restricted to ATCs, with macrophages that can account for up to 57% of the nucleus in ATC, as previously shown (8). The TCGA does not include ATC samples, thus this group is not present in the TCGA correlogram. The third cell type group included epithelial and vascular nuclei (endothelial + pericytes) which could be attributed to the presence of a structured epithelium not perturbed by desmoplasia. Thus, although many high-granularity cell states are idiosyncratic, the proportions of low-granularity cell types still conform to simple statistical trends.

We observed that B and T cells are the dominant cell type in several tumors (Figures 2B, 3A and 3B). Tumor with lymphocyte dominance is confirmed by a reanalysis of the scRNA-seq data from Lu et al., where the fraction of B lymphocytes reached 33% and the fraction of T lymphocytes 65% in one PTC sample (Figure 3C-D) (7). Three PTCs were composed of more than 15% of B lymphocytes and 20% of T lymphocytes in our dataset (Figure 3A-B). At least one of these is known to have a thyroiditis background. Samples with high B cell content also have a large T cell compartment (Spearman’s ρ=0.73, Figure 2C). Do T and B cells colocalize and thus potentially interact via short range communication? Our spRNA-seq data revealed an extra level of idiosyncrasy in those three high-inflammation PTCs. They all displayed sharply circumscribed immune foci (Figure 3C), but the content of these varied. Foci in PTC3 and PTC6 contained both T and B lymphocytes which were also excluded from the epithelial areas. In PTC7 the foci only contained B lymphocytes while T lymphocytes colocalized with the epithelial areas (Figure 3F-G, Supplemental Figure 3A). Thus B and T lymphocytes colocalized in two out of three high-inflammation samples.

**Figure 3.**
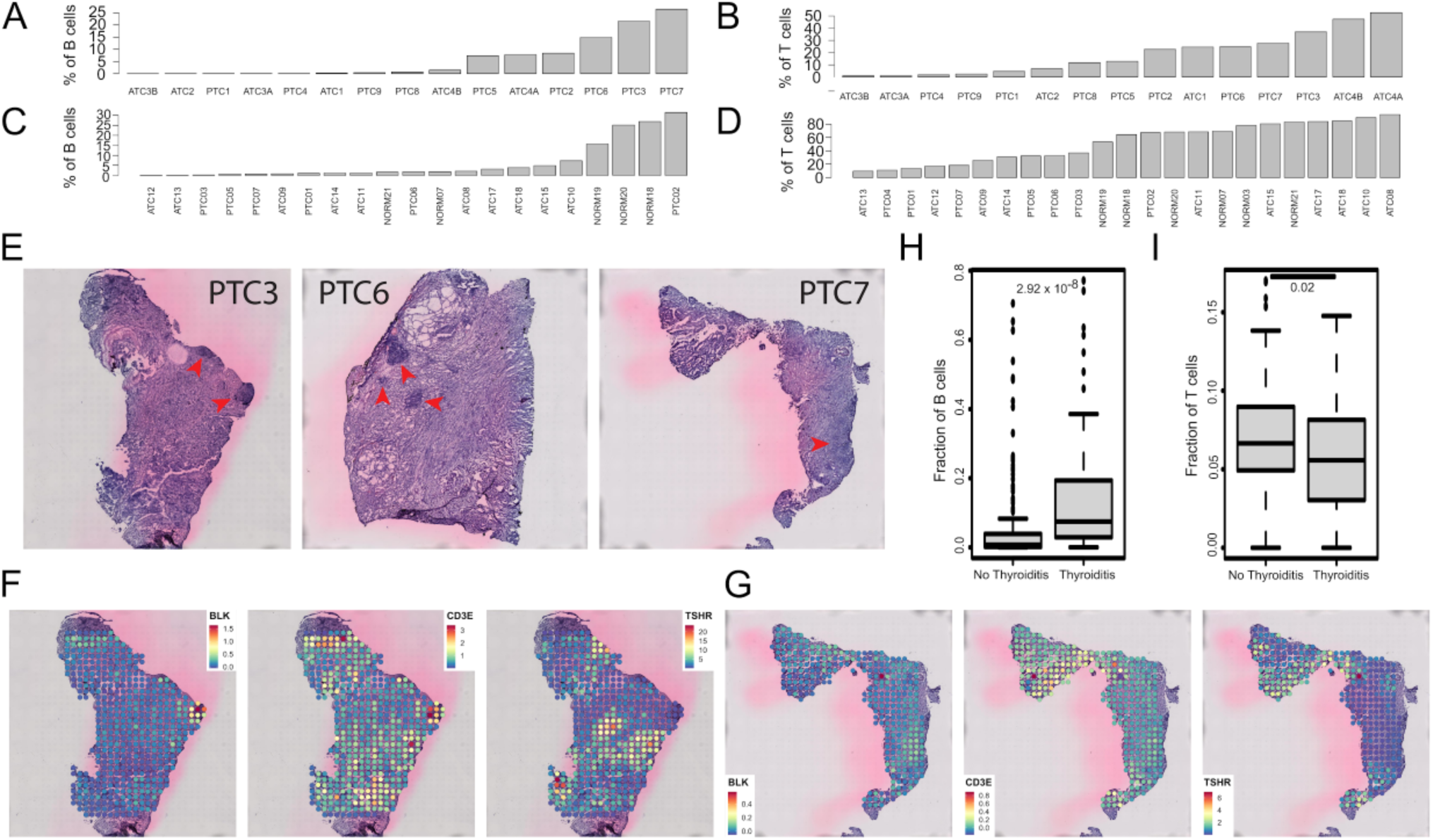
Distribution of B and T cells compartments. **(A)** Fraction of B lymphocytes per sample in our snRNA-seq dataset. **(B)** Fraction of T lymphocytes. **(C)** Fraction of B lymphocytes per sample in data from Lu et al. **(D)** Fraction of T lymphocytes per sample in data from Lu et al. **(E)** H&E images of the 3 PTC samples with the largest B cell fraction. Red arrows highlight immune foci. Expression level B lymphocyte marker BLK, T lymphocyte marker CD3E and thyroid marker TSHR in sample PTC3S3 **(F)** and PTC7S1 (**G**). Cibersortx-inferred fraction of B H**F**) and T lymphocytes (**I**) in 234 *BRAF^V600E^* PTC samples from TCGA with and without thyroiditis background.

Inference of cell type proportions in TCGA (Supplemental Figure 3B) confirmed B cell enrichment in several subsets of *BRAF^V600E^*-mutated PTCs. Furthermore, the B cell fraction was significantly positively associated with patients with thyroiditis background (Figure 3H : median 0.8% vs 7.4%, Wilcoxon rank test *p* = 2.29×10^-8^). By contrast, T cell fraction was negatively associated with thyroiditis background (Figure 3I; median 6.6% vs 5.6%, Wilcoxon rank test *p* = 0,02). In conclusion, B and T cells are major components in a substantial number of PTCs, but do not necessarily colocalize. B cells are associated with a thyroiditis background.

The mean myeloid content in ATC samples was 19.6% (Supplemental Figure 4A). A clear-cut dichotomy between PTCs and ATCs was visible in the myeloid sub-clustering presented in Supplemental Figure 4B. Eighty five percent of PTC myeloid cells belonged to the cluster containing the most cells in the center of the UMAP. The second and third largest clusters were patient-specific (ATC1 and ATC2) and the fourth cluster by size was composed of macrophages from two ATC patients (ATC3A-B and ATC4A-B).

The top marker distinguishing ATC1 myeloid nuclei from all other tumors’ myeloid nuclei was *HIF1A*, a marker of hypoxia (Supplemental Figure 4C). Gene set enrichment analysis shows that ATC1 nuclei are enriched for hypoxia pathways (Supplemental Figure 4D), and comparison of ATC1 and ATC2 nuclei reveals an overexpression of *HIF1A* in most ATC1 cell types (Supplemental Figure 4E), suggesting that hypoxia affects the whole sample.

Thus, thyroiditis, hypoxia and possibly other tumor-wide features, contribute to idiosyncratic expression across different cell types.

### Canonical follicular cell markers are lost in cancer in the exact reverse-order to thyroid organoid differentiation, suggesting a novel classification of epithelial cell states in PTCs and ATCs

We observed a wide range of thyrocyte dedifferentiation states, from fully functional hormone-producing nuclei (Epith *TSHR^+^ TPO^+^* on Figure 4A) to completely dedifferentiated nuclei with mesenchymal features (Epith *TSHR*^-^ on Figure 4A, see also Supplemental Figure 7). In between, we observe clusters at several stages of dedifferentiation that we classified using the four canonical thyrocyte markers involved in thyroid function and homeostasis: the sodium iodide symporter (*SLC5A5*, a.k.a. *NIS*), the thyroid peroxidase (*TPO*), the thyroglobulin (*TG*) and the thyroid stimulating hormone receptor (*TSHR*).

**Figure 4.**
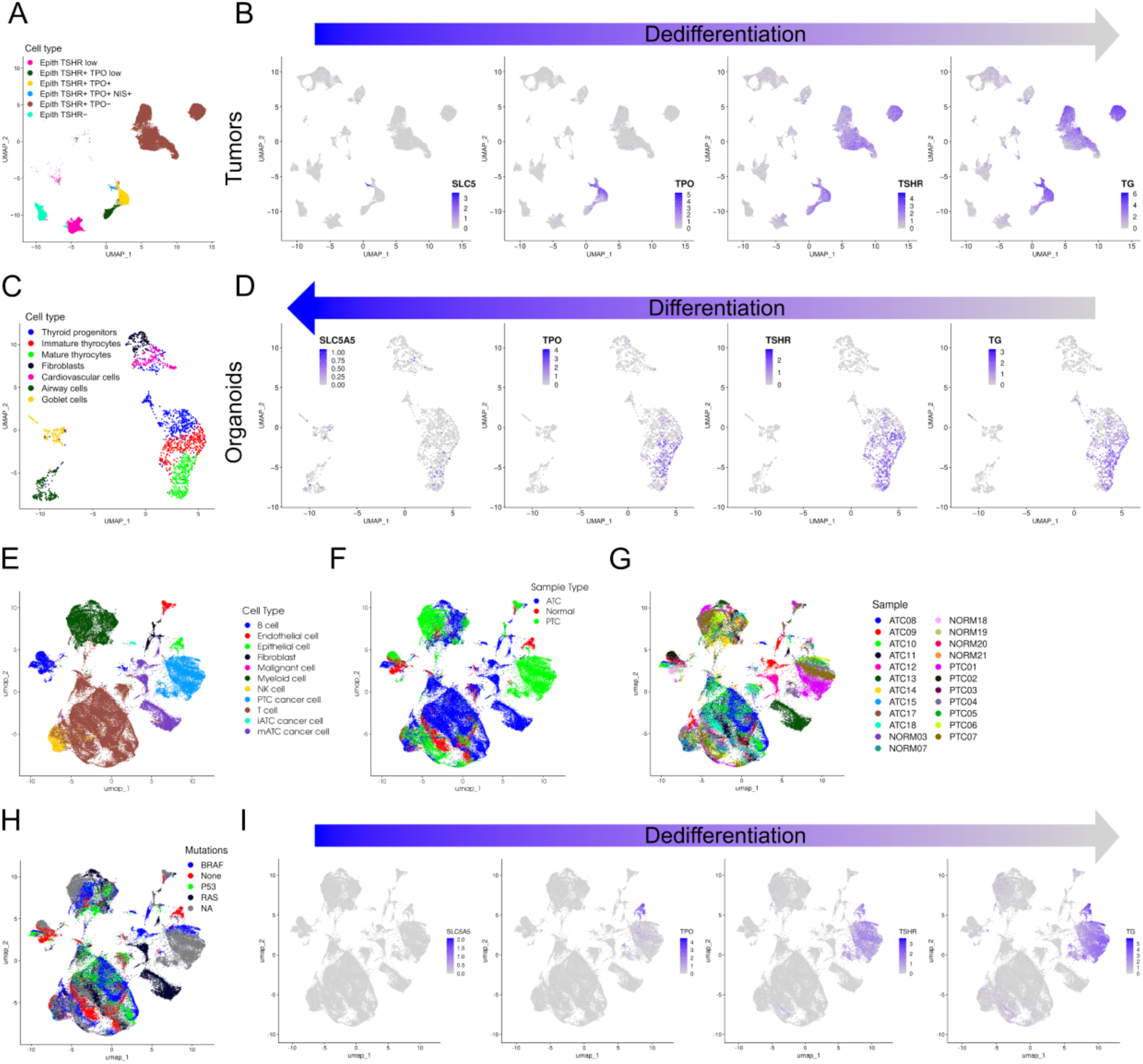
Differentiation and dedifferentiation in development and cancer. **(A)** Cell type annotation for nuclei of epithelial origin in 9 PTC and 6 ATC samples. **(B)** Expression of canonical thyroid differentiation markers in 9 PTC and 6 ATC samples. **(C, D)** Same as A and B for human organoids. Tumor progression is a dedifferentiation process, while organoid development is a differentiation process (arrows). Remarkably, cell subsets are related by the same inclusion relationship, SLC5A5 ⊂ TPO ⊂ TSHR ⊂ TG, in both systems. **(E-H)** UMAP of data from Lu et al. with 6 normal, 7 PTC and 10 ATC samples with cell type annotation as supplied by the authors **(E)**, sample type annotation **(F)**, sample annotation **(G)**, and driver mutation annotation **(H)**. “None” values mean that BRAF, RAS and TP53 were reported as wild-type by the authors. “NA” values were assigned to tumors whose mutational status was not reported by the authors. **(I)** Expression of canonical thyroid differentiation markers in data from Lu et al. with 6 normal, 7 PTC and 10 ATC samples.

Importantly, there was a subset inclusion relationship between these markers with a gradual loss of *TPO*, then *TG* and finally *TSHR* (Figure 4B). This gradual loss could also be observed in the scRNA-seq dataset by Lu et al. (Figure 4I) that includes ATC and PTC tumors harboring the BRAF^V600E^ mutation (Figure 4F-H) (7). Some nuclei in our dataset with detectable levels of the three aforementioned markers also had a detectable expression of *NIS* (Figure 4B). Hypothesizing that this inclusion relationship could be a fundamental feature of the thyrocyte (de)differentiation process, we measured the same four markers in a recent human scRNA-seq thyroid organoid dataset (28) (Figure 4C). It revealed a gradual gain in the thyroid progenitors, immature thyrocytes and mature thyrocytes clusters of *TSHR* (15%, 62%, 78%), then *TG (*21%, 60%, 79%), *TPO* (11%, 30%, 55%) and *NIS* (1%, 2%, 10%) (Figure 4D).

Thus, the ordering of thyrocyte differentiation marker acquisition in thyroid development is mirrored by their loss in cancer, suggesting a common differentiation and dedifferentiation time-reversible axis. We propose to classify epithelial nuclei according to this fundamental structure. We identified six cell categories spanning the thyrocyte differentiation spectrum in our dataset: Epith *TSHR^+^ TPO^+^ NIS^+^* nuclei (*NIS*^+^), Epith *TSHR^+^ TPO^+^* nuclei (*TPO*^+^), Epith *TSHR^+^ TPO^low^* nuclei (*TPO^low^*), Epith *TSHR^+^ TPO^-^*(*TPO*^-^) nuclei spanning most PTC samples, and finally Epith *TSHR^low^* nuclei and Epith *TSHR^-^* nuclei that are solely coming from ATC samples.

### TSHR^low/-^ cells have undergone full-blown EMT and are found in ATCs

We demonstrate through specific snRNA-seq experiments mixing cells from tumors and adjacent healthy tissue that TPO^+^ cell are healthy, non cancer, cells (Supplemental Figure 5) *TSHR^low^* and *TSHR*^-^ nuclei were ATC-specific and idiosyncratic, 95.2% originated from ATC2 and 94.5% from ATC1, respectively. *TSHR*^-^ nuclei are fully dedifferentiated, they do not express any epithelial markers (Figure 4B and Supplemental Figure 7B) and express numerous mesenchymal markers, including extracellular matrix components and mesenchymal transcription factors that are also present in *bona fide* fibroblasts (Supplemental Figure 7B). Their malignant status is confirmed by the presence of chromosomal aberrations detected by inferCNV (29) (Supplemental Figure 6A). *TSHR^low^* nuclei are at an intermediate stage of dedifferentiation : they express—at lower levels—many of the aforementioned mesenchymal markers (Supplemental Figure 7A), and lost most epithelial markers (Figure 4B and Supplemental Figure 7B), but not all : 26% of *TSHR^low^*nuclei still express *TSHR* (Supplemental Figure 7B). The top 10 differentially expressed genes between *TSHR^low^* and *TPO*^+^ nuclei are mesenchymal markers that are also expressed in fibroblasts, but not in PTC-specific subsets (Supplemental Figure 7B). Furthermore, motif analysis with SCENIC (30) suggests a role of ZEB1 in driving ATC cell EMT (Supplemental Table 9).

The percentage of nuclei expressing the proliferation marker *TOP2A* is depicted in Supplemental Figure 7C. *TSHR^low/-^*cells were the fastest proliferating compartment of epithelial origin (6 to 14% *TOP2A*^+^ nuclei), along with a cluster of cells with high mitochondrial activity originating at 95% from ATCs. Conversely, PTC epithelial nuclei proliferated slowly (less than 0.4% *TOP2A*^+^ nuclei). Taken together, these results show that *TSHR^low/-^* ATC cells combine aggressiveness features including full-blown EMT and high proliferation.

Beside ATC1 and 2, two remaining ATC sample pairs (ATC3A-B and ATC4A-B) came from 2 patients. The cell type proportions within each pair were more comparable. Little epithelial or EMT cell types were present in these samples (Figure 2B). This could be explained by low purity: 86.8% and 81% of nuclei belong to the microenvironment in ATC4A and ATC4B, respectively. Fifty eight and 84.7% of nuclei in ATC3A and ATC3B belong to the high mitochondrial gene expression cluster.

High expression of mitochondrial genes typically result from technical factors or cell death. ATC3A-B had a RIN above 7 and all nuclei included in the dataset passed the quality control thresholds for read counts and number of features. Caspase 3 immunofluorescence stainings, on the other hand, revealed numerous clusters of epithelial cells at the terminal stages of apoptosis inside the lumens of epithelial structures (Supplemental Figure 7D). An intraluminal location of apoptotic thyrocytes has been previously reported (31, 32). A sub-clustering of this cluster reveals various cell types, both from the microenvironment and of epithelial origin. Therefore cell death is systemic in those samples and the relative absence of nuclei of epithelial origin in ATC3 and ATC4 can be explained by their high abundance in the high mitochondrial expression cluster, which also has the second highest proliferation rate (Supplemental Figure 7C). SpRNA-seq further revealed that in ATC3B (Supplemental Figure 7E) a crisp border delimits the area with high mitochondrial gene expression and thus the necrotic region. Most genes are overexpressed on the other side of this border as exemplified in Supplemental Figure 7E with expression of macrophage marker *CD68*.

### TSHR^+^/TPO^-^ cells are a hallmark of PTC, their highly idiosyncratic states are partly associated with DNA copy number and morphological variations

The *TPO*^-^ cluster was composed of 30,609 nuclei originating exclusively from PTCs in which it was the dominant cell type in most cases (Figure 2B). Global clustering and subclustering of this cell type revealed one generic and three idiosyncratic clusters for PTC4, PTC9, and part of PTC2 (Figure 1A).

Some of these can be explained by chromosomal aberrations. InferCNV (29) analysis shows that the idiosyncratic cluster containing all PTC4 epithelial nuclei harbors a deletion on chromosome 13 (Supplemental Figure 6B).

PTC2 epithelial nuclei are distributed among the generic *TPO*^-^ cluster and a PTC2-specific cluster. This clear-cut separation was mirrored in the inferCNV results which revealed a diploid clone and a clone with gains on chromosomes 3, 7 and 8 and a deletion on chromosome 22 (Figure 5A). The latter is a recurrent CNV present in 10% of PTC samples from TCGA (11). The aneuploid cluster under-expressed most thyroid differentiation markers, including *TG* (Wilcoxon p-value = 2.2 × 10^-16^, Figure 5B). The expression of *TG* and the median expression of genes located on the aberrant chromosomes reveal a crisp spatial boundary between the euploid and aneuploid regions on all 6 slices of the PTC2 sample (Figure 5C). A closer examination of these 2 regions reveals a subtle morphological difference in their tissue structure. The 2D sections of epithelial sheets run continuously over distances of several millimeters in the euploid region. By contrast, they run over distances of a few hundreds micrometers in the aneuploid region (Figure 5D). This subtle difference, which initially escaped our attention, suggested that aneuploidy results in a change in the tissue’s mechanical properties.

**Figure 5.**
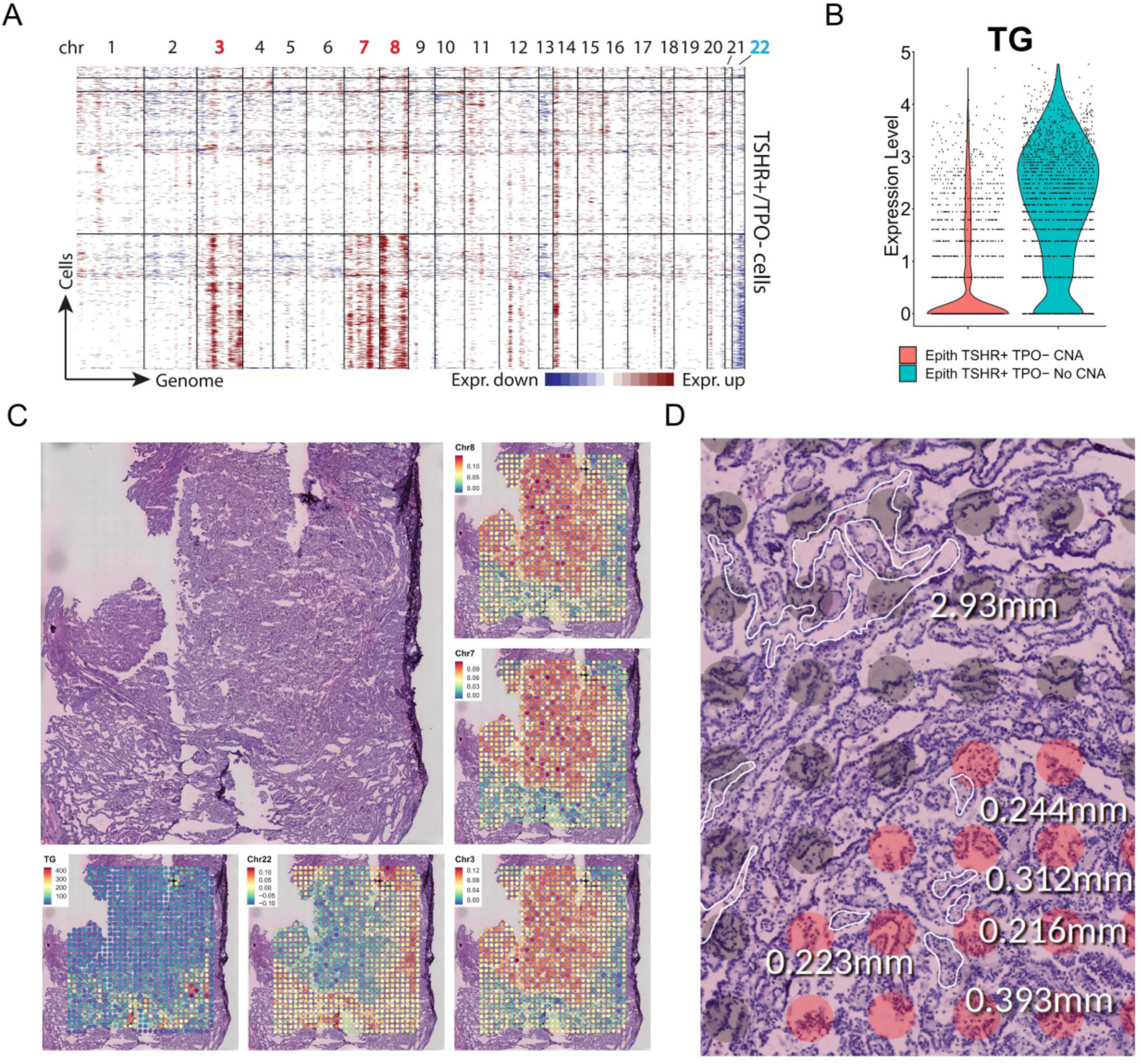
DNA copy numbers and morphology in PTC. **(A)** DNA copy number alterations inferred from snRNA-seq by infercnv in sample PTC2. **(B)** Expression level of *TG* in nuclei with and without chromosomal aberrations in PTC2. **(C)** H&E image of PTC2 and expression levels *TG* and genes present on aberrant chromosomes. **(D)** H&E image showing morphological differences between the area bearing chromosomal aberrations (red dots) and the rest (gray dots). The length of epithelial sheets’ sections is shorter in the CNA area.

No copy-number changes were found in PTC9. High-depth targeted sequencing of a panel of 380 cancer genes revealed no additional mutation beyond the canonical *BRAF^V600E^*. Thus, the idiosyncratic expression of this tumor remains unexplained.

### Mesenchymal features, including Fibronectin 1 production, of TSHR^+^/TPO^-^ cells are highly variable within individual PTCs

Idiosyncratic expression restricts integrative methods such as trajectory analysis across samples. We instead structured our investigation of *TSHR*^+^/*TPO*^-^ cells on an established generic feature of *BRAF^V600E^* PTC. *FN1* has been associated with *BRAF^V600E^* PTC in previous gene expression studies (33). More specifically, a reanalysis of TCGA (11) shows that it is a hallmark of *BRAF^V600E^* PTC, with a 100-fold overexpression (Figure 6A).

**Figure 6.**
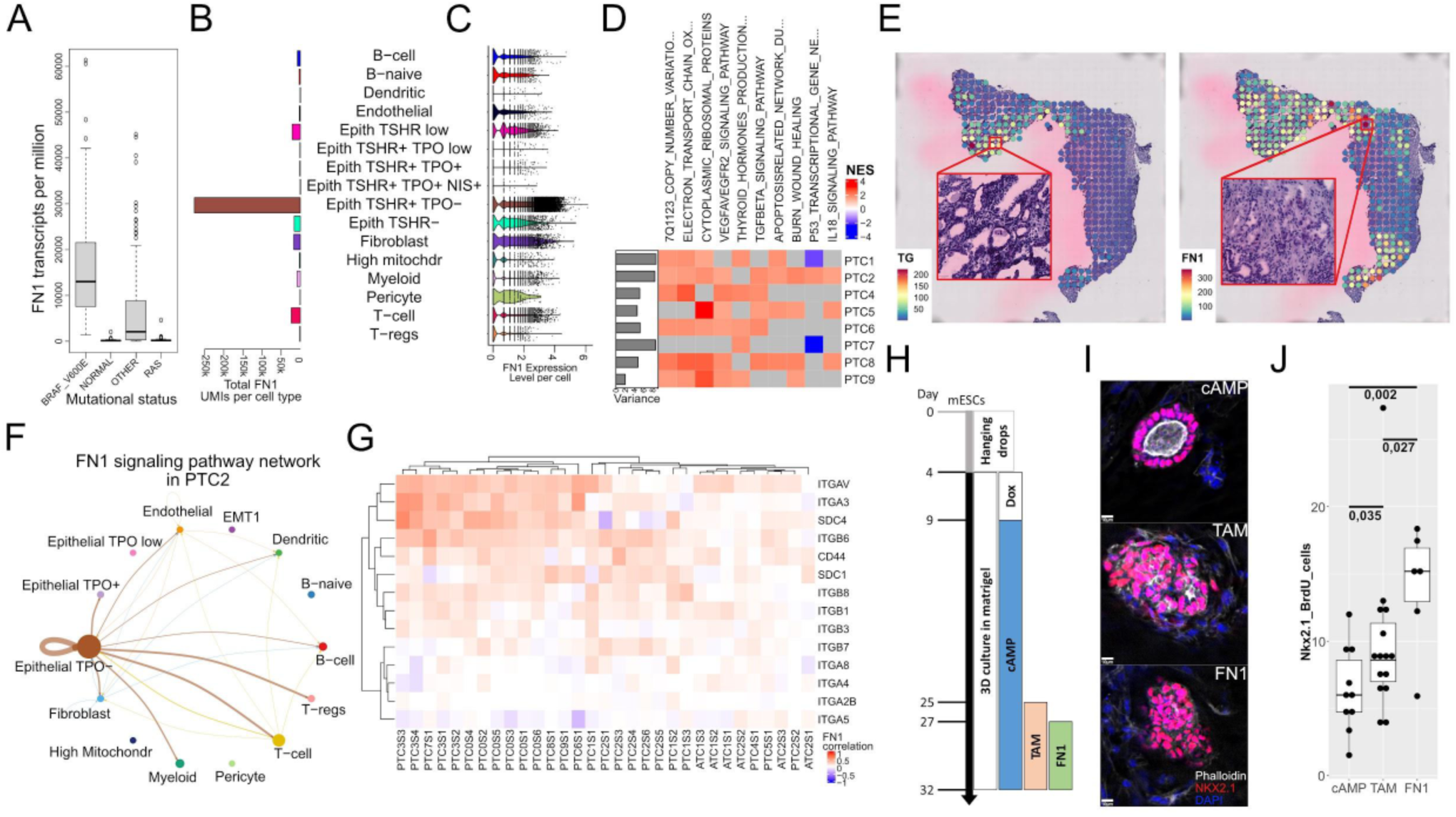
Mesenchymal features in PTC. **(A)** Expression of *FN1* in the 568 TCGA samples. **(B)** Total amount of *FN1* UMIs per cell type in 9 PTC and 6 ATC samples. **(C)** Per cell expression of *FN1* across cell types. **(D)** Reactome pathways significantly associated with the top 3 components of transcriptional variation in *TSHR^+^*/*TPO*^-^ cells. The proportion of variance explained by the components is displayed in the left-side barplot. **(E)** Expression of *TG* and *FN1* in PTC7S1 with zoom on high *TG*/low *FN1* (left) and low *TG*/high *FN1* (right) areas. **(F)** Ligand-receptor interactions for the FN1 signaling pathway in sample PTC2 as inferred by CellChat. **(G)** Correlation of *FN1* expression level with *FN1* receptors’ expression level across spots in spatial transcriptomics samples. **(H)** Timeline of the thyroid cancer organoid differentiation protocol. Doxycycline for *Nkx2.1*-*Pax8* induction and Tamoxifen for *Braf^V637E^*induction. **(I)** Immunofluorescence stainings for *Nkx2.1* (pink) and phalloidin (white) among the different conditions (images representative of the 3 experimental replicates). **(J)** Percentage of *Nkx2.1* positive cells that are BrdU positive across the 3 experimental conditions.

As a component of the extracellular matrix, *FN1* is expected to be produced by mesenchymal cells. Our data showed, however, that it was mainly produced by epithelial nuclei, both in absolute quantities (Figure 6B) and relative per-nucleus counts (Figure 6C). Individual cancer cells expressed on average 40% more *FN1* than fibroblasts. Its expression in the *TPO*^-^ compartment was negatively correlated with some differentiation markers : Spearman’s ρ was -0.45 between *TG* and *FN1*.

Dedifferentiation is the main source of transcriptional variation for some tumors but not all. To illustrate this we assessed the quantitative contribution to the overall transcriptional variation within the *TPO*^-^compartment of individual tumors. The principal components of transcriptional variation were computed (see Methods), and their associations with Reactome pathways were evaluated. Among the principal sources of variations are pathways associated with differentiation and EMT such as THYROID_HORMONE_PRODUCTION and TGFBETA_PATHWAY and additional pathways related to inflammation, vascularization and metabolism (Figure 6D).

SpRNAseq experiments showed that regions expressing high amounts of *FN1* and low amounts of thyroid differentiation markers were associated with a loss of epithelial structure (PTC7, Figure 6E). In PTC2, aneuploid, dedifferentiated nuclei expressed more *FN1* (Wilcoxon p-value = 2.2 × 10^-16^).

Thus, our results show that EMT is partial even in PTC tumors where dedifferentiation is the main source of transcriptional variation; none of the mesenchymal canonical markers, transcription factors and matrix proteins observed in ATC are expressed as strongly in PTC (Supplemental Figure 7A-B).

### Fibronectin 1 increases proliferation of cancer cells through an autocrine loop

Finally, connectome analysis demonstrates that potential receptors of *FN1* are prominently expressed by *TPO*^-^ nuclei, suggesting an autocrine loop as shown with PTC2 in Figure 6F. Correlations between the expression of *FN1* and its candidate receptors across spots of our spRNA-seq PTC slices are mostly positive as shown by Figure 6G.

This recurrent association across studies, along with the unexpected production by epithelial cells and predicted autocrine mechanism prompted us to test the effects of Fibronectin 1 on thyroid cancer in vitro. Using a mouse ESC-derived thyroid organoids model in which a Braf-oncogene could be induced by tamoxifen treatment (34, 35), we explored the effect of *Fn1* treatment on cell proliferation. This model expresses both *Fn1* and the receptors of *Fn1* signaling. We used uninduced controls, Braf-induced organoids (TAM), and Braf-induced organoids treated with soluble *Fn1* (FN1) (Figure 6H-I). Combining BrdU uptake and Nkx2-1 staining, we observed that after 7 days of treatment, double-positive cells were 1.43 fold higher in cancerous thyrocytes compared to the controls while a 1.77 fold increase was observed between *Fn1*-treated and cancerous thyrocytes (Figure 6J). This indicates that *Fn1* treatment further increases the proliferation of thyroid cancer organoids *in vitro*.

## Discussion

We demonstrated from cell mixing experiments that the transcriptional variations within the same cell types coming from different patients is of biological and not technical origin. This finding may apply to other studies as several idiosyncratic clusters were associated with aneuploidy, a widespread phenomena in cancer. Thus, a large fraction of the transcriptional variation among PTC cells, and most probably cells from other cancers, is driven by patient-specific idiosyncratic factors. Of note, we also found idiosyncratic expression in macrophages and T cells from ATCs, suggesting a diversity of cell states yet to be explored.

Our cell mixing experiments revealed a potential shortcoming in the application batch effect correction algorithms that are widely used in single cell studies. We demonstrate that technical batch effects are minimal in our study, thus these algorithms were not needed. But they would have totally concealed idiosyncratic transcriptional variation, giving the erroneous impression that cells from different patients form an homogeneous group. We could avoid this pitfall thanks to our cell mixing experiments. Most single cell studies, however, do not present such experiments. In these studies, the decision to apply or not batch correction and the selection of batch correction algorithm hyperparameters rest on the typically implicit *a priori* assumption that technical variation dominates over biological variation. But this assumption is rarely backed by experiments. As a result, biologically essential idiosyncratic expression may be grossly underestimated and flawed analysis may follow. For example, the erroneous merging in a single large cluster of epithelial PTC cells from all patients following batch integration in Fig 1B would have called for a cross-patient trajectory analysis, as performed in many studies. It would have been flawed, however, since cells from, say PTC4, have a chr13 deletion and therefore a unique idiosyncratic cell state with no plausible continuous transition from/to other cell states in the dataset.

We propose that the principle of our cell mixing experiment is applicable systematically to ensure that batch correction does not corrupt biological signals. Cell’s patient identities can be recovered computationally from their SNPs when these are known or, alternatively, a sample multiplexing kit can tag them per patient. Thus, cells can be mixed in vitro and their patient identity can be recovered in silico. Once all cells from a study are mixed their transcriptomes may be determined from as many runs as needed to reach the desired cell count. Since, by construction, the same cell populations/clusters are present in all runs (these are also known as anchors), the variation between different runs is necessarily technical and can be removed efficiently and safely by batch correction algorithms. As the cost of single cell technology drops, this approach could enable the profiling and integration of very large series of specimens for which the consistency of experiments is harder to control.

Our snRNA-seq dataset includes 13 thyroid cancer patients. We choose to focus on BRAF-mutated PTC and to ATC samples to keep to a level compatible with the cost of single cell technology the number of disease categories to be addressed. Unfortunately, the ATC samples included in this study do not have BRAF mutations, which limits comparisons between both tumor types. This only paints a restricted picture of the many facets of the disease and their heterogeneity in the patient population—much of which remains to be discovered. As such, a complete stratification scheme cannot be established from our small cohort. Furthermore, there is a real possibility that some features that appear as idiosyncratic in this study will be shared within tumor subgroups that may be revealed in future large scale studies. This work should be viewed as a stepstone for these studies. It should also be noted that the 13 patients or the 18 samples taken from them, even though typical for a single cell study, constitutes a very limited number of biological replicates. As such, it limits the amount of clinical correlates that can be computed with sufficient statistical power. As explained in further details by Squair et al. (36), the illusion of statistical power provided by the thousands of cells captured in a single cell dataset quickly fades when taking into account the actual number of biological replicates that are the relevant entities when computing clinical correlates (36).

The first microenvironment idiosyncrasy evidenced here is the abundance of T and especially B lymphocytes that is associated with thyroiditis in the TCGA. Reports about the abundance of B cells in PTC are conflicting, suggesting either an enrichment (37) or a depletion (38). Estimation of B cell proportions from TCGA data hints at the reason for these inconsistencies, as they vary widely depending on the subcategory of PTC considered. Previous studies suggest an increased risk of developing thyroid cancer for thyroiditis patients (39, 40) but the exact mechanisms underlying that transition need to be further studied, including the role played by infiltrating immune lymphocytes at the onset and during the progression of the disease.

The second microenvironment idiosyncrasy concerns myeloid cells and is especially marked in ATC samples. ATC cells in general display a greater inter-patient heterogeneity for most cell types, hinting at a greater ability of fully dedifferentiated cancer cells to remodel not only themselves but also their microenvironment. Our attempts to classify macrophages in discrete and well defined categories as described in the literature (41, 42) failed. This could result from a technical limitation of our dataset. For example, key marker genes expressed in the cytoplasm may be invisible in our single nuclei profiling. Alternatively, these inconsistencies could reflect limitations of those over-simplified discrete classifications of a cell type that is extremely complex, labile and that possibly varies within a continuum (43). The only discernible patterns to explain those idiosyncrasies were mostly cell states (e.g. hypoxia, metabolic activity, etc.). Their consequences on the disease prognosis need to be further studied.

This dataset encompasses the full spectrum of thyroid cancer dedifferentiation, from fully differentiated normal hormone-producing thyrocytes to fully dedifferentiated mesenchymal cancer cells. We defined the main stages of this dedifferentiation process based on the expression levels of four canonical thyroid markers (*TSHR*, *TPO*, *TG* and *NIS*). Other thyroid markers present the same profile as some of these four canonical markers, including transcription factors. But this selection has the advantage of being composed of well established thyroid markers that are strongly expressed (except for *NIS*), making them less susceptible to dropouts. The technical issues discussed hereafter surrounding *NIS* render it dispensable for this classification, but we chose to retain it because of its peculiar expression profile in our dataset and prominent role in thyroid biology. Remarkably, the loss of these markers with dedifferentiation in cancer mirrors their gain during thyroid organoid development, suggesting a fundamental relationship between these two phenomenons. These markers accurately matched the clustering based on whole transcriptomes, suggesting their ability to summarize global cell states. They proved to be useful to organize the diverse array of transcriptional cancer cell states in this study. These four markers have already been established as prognostic markers in thyroid cancer by various publications over the years (44–48). They are also part of the TDS and thyroid index, two scores with prognostic value that are based on bulk gene expression data (11, 49). But as the TDS and thyroid index operate on bulk transcriptomes, they average the expression profiles of numerous cancer and non-cancer cells. Thus, they not only measure dedifferentiation but also other factors like purity (Supplemental Figure 8). By contrast, our stratification scheme applies to cells and not tumors. It measures dedifferentiation directly and provides a view of the intra-tumoral heterogeneity.

Fully differentiated and functional thyrocytes express *TSHR*, *TPO* and some of them *NIS* according to snRNA-seq data. The organoids from Romitti et al., albeit functional, also lack a detectable expression of *NIS* in single cell RNA-seq (Figure 4D), while its expression is detectable in bulk RNA-seq (28). Other PTC studies have also shown that the mRNA and protein expression levels of NIS, or even the various subcellular localizations of the NIS protein, do not correlate equally with clinical characteristics (48, 50), further highlighting the technical challenges surrounding NIS. This suggests that the lack of *NIS* expression in our functional thyrocytes profiled with snRNA-seq might be attributable to dropouts rather than true lack of expression. In those nuclei the expression level of *NIS* might be below the detection threshold of snRNA-seq. This observation can also be true for the other markers used here. For instance, TPO negative cells don’t pass the threshold for snRNA-seq detection, but they might still express TPO mRNA at lower levels. One patient with Hashimoto thyroiditis expresses *NIS* at levels above its detection threshold. NIS expression is supposedly increased by Graves’ disease and decreased by Hashimoto thyroiditis (48, 51–53). It is thus surprising that normal adjacent cells with Hashimoto background express more NIS than the other normal adjacent cells. To decipher this, one would need to establish a comprehensive single cell RNA-seq profile of normal thyrocytes in healthy individuals and compare it to thyroids in different pathological settings. As these lines are written, the only publicly available single cell RNA sequencing dataset available for healthy human thyroids is the human cell landscape (54). This thyroid dataset originates from 2 individuals and one of them has thyroiditis. Both have low numbers of UMI per cell.

Fully dedifferentiated cancer cells do not express any of the aforementioned canonical markers. Our dataset includes one such fully dedifferentiated ATC and another one almost fully dedifferentiated that keeps a residual expression of some epithelial markers mixed with mesenchymal ones (dubbed *TSHR^low^*). These two clusters represent the only cancer cells in our dataset that undergo drastic EMT and they only include ATC cells, not PTC. Previous reports mentioned signs of EMT in PTC (55) but our dataset and other accumulating PTC datasets show that it is only partial and heterogeneous in individual tumors (5–7, 17, 23, 24, 56).

The cancer epithelial cells from our *BRAF^V600E^*PTC samples are characterized by a loss of *TPO* and *NIS* expression but neither *TSHR*, nor *TG*. They form one main cluster segregated from normal and ATC cancer cells. Yet, this main cluster displays varying degrees of differentiation between tumors and also within individual tumors. This differentiation continuum can be observed in thyroid organoids as well but the link between this differentiation level and thyroid function remains to be elucidated. In addition to differentiation level, idiosyncrasies pertaining to metabolic activity or other cell states have also been observed. One of the most marked sources of heterogeneity in this cluster is chromosomal aberrations. Among those chromosomal aberrations are deletions of chromosome 12 and 22 that were already documented in thyroid cancer literature (11, 57). Interestingly, the deletion of chromosomal arm 22q has been associated with differentiated thyroid cancer co-occurring with ATC (57). This is especially relevant considering that the cancer cells bearing that deletion in our PTC2 sample have a more mesenchymal phenotype and are less differentiated than other cancer cells in the same sample. This specific example highlights the potential of combining snRNA-seq and spRNA-seq to establish that link between tissue morphology and molecular profile (and clinical characteristics), despite the complex nature of this heterogeneous tumor with subclonal cancer populations. Linking an objective assessment of tissue morphology to molecular profiles and clinical outcomes in a systematic way at the subclonal level would enable routine imagery techniques used in pathology departments to bring much more valuable information.

Several articles studying ATC and PTC have established a link between cancer cells originating from both using trajectory inference (7, 17). Lu et al. describe a population of inflammatory PTC cells at the intersection between both tumor types. We have observed no such intermediate state cluster in our 9 PTC samples. Even though trajectory analysis would certainly find an expression gradient between our ATC and PTC cancer cells, we have no reason to believe there should be a direct connection between them and thus this gradient would bear no biological meaning. The absence of such intermediate states is likely a limitation of our dataset that could be explained by two factors. The first is that it includes a relatively small number of samples, like most single cell studies, thus limiting the probability to capture every intermediate stage of the disease categories we selected. The second is that our PTC samples have the BRAF mutation while our ATC samples unfortunately do not. However, it should be noted that trajectory analysis algorithms only measure gradients in the expression data (58). As such they are not concrete evidence of filiation between the cell populations like lineage tracing experiments are.

As explained above the differentiation level of *BRAF^V600E^* mutated cells varies between PTCs and within individual PTCs. The loss of differentiation markers is correlated to the acquisition of a mesenchymal phenotype in those cells. Fibronectin 1 is a mesenchymal marker that is overexpressed in BRAF mutated PTC and is inversely correlated to thyroid differentiation markers. It has been associated with increased migration, invasion and proliferation in thyroid cancer (59–62) but the cell type responsible for most of its production in human thyroid cancer hasn’t been established in a quantitative manner. Our results demonstrate that it is the cancer cells themselves that produce FN1 and its integrin receptors, thus forming an autocrine loop. Several types of analysis associate it with the TGF signaling pathway and inflammation but the exact mechanisms triggered in human thyroid cancer cells need to be further studied. Its effect on proliferation in vitro might be explained by an interaction with the MAP kinase and/or PI3 kinase pathways and some studies also associate it with resistance to kinase inhibitor treatments in thyroid cancer and melanoma (62, 63). However, our results in vitro do not allow us to disambiguate between an interaction with the oncogenic pathway and a more generic effect on thyroid epithelial cells proliferation, and more experiments using this model would be required to do so.

In conclusion this dataset offers insights that could help develop new therapeutic strategies, improve risk stratification for thyroid cancer patients and better understand the disease. We confirmed previously published findings and highlighted potentially relevant new avenues, but most of all we show the potential of combining single-cell resolution RNA sequencing with tissue imaging to better grasp the many facets of this highly heterogeneous disease. Some findings need to be further studied using different models. For instance, a human thyroid organoid model where the level of BRAF oncogene activation would be fine tunable could suggest whether the stratification we observe here is a fundamental feature of the BRAF mutated thyrocytes’ dedifferentiation process, or whether it results from interactions with the microenvironment as well. Other findings need to be confirmed with larger scale studies that could build upon the technical lessons learned here to make the best possible experimental designs and get a complete picture of thyroid cancer.

## Methods

### Tumor tissue samples

OCT-embedded tumor tissue samples were obtained from Institut Jules Bordet and Centre Hospitalier Universitaire de Liège’s biobanks. This study was approved by the local ethics committee (protocol #1978). After surgical resection, tumor tissue samples were cut with a disposable scalpel (Swann-Morton, 0510) to fit the Tissue-Tek® Intermediate Cryomold® (Sakura, 4566). Tumors were embedded in O.C.T (Leica, 14020108926), freezed at -80°C and stored at -80°C.

All samples selected had an RNA integrity number (RIN) value of 7 or higher. Only PTC samples harboring the *BRAF^V600E^*mutation were selected. RIN values and BRAF mutational status were determined as previously described (64).

### Parallel processing of tumor samples

O.C.T. embedded tumor tissues were sliced using a UV cryostat (Leica, CM1950). All slices for immunofluorescence (IF) and spatial transcriptomics (spRNA-seq) experiments were cut at the same time to maximize their likeness. Each spRNA-seq slice was 10µm thick and surrounded by 3 IF slices on both sides that were each 6µm thick. spRNA-seq slices were transferred on library preparation spatial transcriptomics slides and IF slices were transferred on SuperFrost® Plus microscope slides (Thermo scientific, j1800amnz). Slices used for single nuclei sequencing and genotyping were 20µm-thick and collected in Eppendorf® DNA LoBind tubes (Eppendorf, 0030108051). The number of slices per snRNA-seq or genotyping experiment ranged from 5 to 15 depending on the sample size. All slides and tubes were stored at -80°C until further use.

### Immunofluorescence

O.C.T. embedded tissue slices of 6µm thickness on SuperFrost® Plus microscope slides (Thermo scientific, j1800amnz) were fixed with 4% paraformaldehyde for 10 minutes at room temperature, washed three times in phosphate buffered saline (Gibco, 14190094) and blocked with PBS containing 1% BSA (Biowest, P6155), 5% horse serum (Invitrogen, 16050130) and 0.2% Triton-X100 (Bio-rad, #1610407) for 1 hour. Primary antibodies were diluted in blocking solution and incubated overnight at 4°C. After 3 PBS washes, secondary antibodies were diluted in blocking solution with DAPI (Sigma Aldrich, D9564) and incubated 1 hour at room temperature. SuperFrost® slides were finally washed 3 times with PBS and mounted using Glycergel Mounting Medium (Agilent, C0563) and cover glass (VWR, 631-0137). Imaging was performed using a Vectra Polaris scanner (PerkinElmer, P/N CLS147553) and images were generated using QuPath v0.4.1 (65). The reference of the antibodies used are provided in Supplemental Table 8.

### Targeted DNA sequencing panel

O.C.T. embedded slices were put in a 1mL WHEATON® Dounce Tissue Grinder (DWK Life Sciences GmbH 357538) containing 500µL of Dulbecco’s phosphate-buffered saline (Gibco™, 14040117). The mix was homogenized and transferred to a new DNA LoBind tube after adding 500µL of DPBS. The tubes were centrifuged 3 minutes at 6000g and the supernatant was discarded. The pellet was resuspended in the proteinase K solution from the DNeasy® Blood & Tissue kit (Qiagen, 69504) and incubated at 56°C overnight. The rest of the DNA extraction was performed according to the kit’s instructions. The extracted DNA was used to construct libraries with the “solid tumors and haematological tumors (STHT)” gene panel version 3 containing 380 genes (BRIGHTCore, Brussels, Belgium). Libraries were sequenced using an Illumina NovaSeq 6000 system.

### spRNA-seq experiments

spRNA-seq was performed using an adapted version of the original protocol by Salmén et al. (66). The tissue was incubated in the eosin solution for 60 seconds, Pepsin/HCl permeabilization was performed during 7 minutes and the two-step tissue removal using 3X β-mercaptoethanol was applied. The glycerol used as a mounting medium prior to image acquisition was pre-chilled at 4°C. Libraries were sequenced using an Illumina NovaSeq 6000 system. Bright field imaging after H&E staining was done using a Nanozoomer-SQ digital slide scanner (Hamamatsu, C13140-01). Fluorescence imaging after Cy3 staining was done using a Genepix 4000B microarray scanner (Molecular devices, GENEPIX 4000B-U).

### snRNA-seq experiments

O.C.T. embedded slices were homogenized in a single nuclei suspension using a modified version of the nucleus isolation workflow for ST-based buffers of Slyper et al. (67). All reagents and buffers used were as defined by Slyper et al.

1mL of cold TST was transferred to a pre-chilled 1mL WHEATON® Dounce Tissue Grinder (DWK Life Sciences GmbH 357538). The 20µm-thick OCT-embedded tissue slices were put at the bottom of the douncer in the TST. The loose pestle was used to stroke 5 times, then the tight pestle 5-10 times or until the solution was homogeneous. The homogenized solution was then filtered through a 40µm VWR cell strainer (10199–655). An additional 1 mL of the detergent buffer solution was used to wash the well and filter. The volume was brought up to 5 mL with 3 mL of 1X ST buffer. The sample was then transferred to a 15 mL conical tube and centrifuged at 4°C for 5 minutes at 500 g in a swinging bucket centrifuge (acceleration max, deceleration 5). The pellet was resuspended in 300µL of 1X ST buffer. The nucleus solution was then filtered through a 30 µm MACS smartstrainer (Corning, catalog no. 352235). Nuclei were counted using a LUNA-FL™ Dual Fluorescence Cell Counter (Logos bioscience, L20001) in bright field counting mode.

Nuclei suspended in ST buffer were diluted to achieve a targeted recovery of 8000 nuclei per sample as specified in the Chromium Single Cell 3ʹ Reagent Kits v3 instructions (10X Genomics, CG000204 Rev D). All further steps of the library preparation protocol were done according to the kit’s instructions. Libraries were sequenced using an Illumina NovaSeq 6000 system following 10X Genomics’ kit instructions.

### spRNAseq processing pipeline

Image files produced by the Nanozoomer-SQ were transformed using ndpi2tiff v1.8 (68), cropped using tifffastcrop v1.3.10 (69), rotated, saved in pyramidal tiff format and resized jpeg using vips v8.4.5 (70). Image alignment and detection of spatial spots was performed using the ST Spot Detector v2.0.2 as described by Salmén et al. (66).

Raw sequencing data was aligned, annotated, demultiplexed and filtered using the spatial transcriptomics pipeline (v1.7.6) developed by Salmén et al. (66) using default parameters with GRCh38 reference genome and gene annotation Genecode v28.

Analyses were done using R 4.1.0 and Seurat version 4.0.3 (71). UMI counts generated by the spatial transcriptomics pipeline, jpeg brightfield images and corrected spot coordinates were used to create a spatial Seurat object using a custom script. The resulting library was scaled and normalized using the SCTransform function from Seurat. Other processing steps were done according to Seurat’s recommendations.

For visualization purposes counts were smoothed in R 4.1.0 with a simple singular value decomposition denoising : ‘svd <- svd(log10(1 + c)); imputed <- svd$u[, 1:10] %*% diag(svd$d[1:10]) %*% t(svd$v[,1:10]); imputed <- 10 ^ imputed - 1’, where c is the raw UMI counts matrix.

### snRNAseq processing pipeline

Raw sequencing data was aligned, annotated, demultiplexed and filtered using Cell Ranger Software (v.6.0.1) with GRCh38 reference and gene annotation Ensembl 98. Analyses were done using R 4.1.0 and Seurat version 4.0.3 (71). Briefly, raw counts from Cell Ranger were loaded and the “background soup” was removed using SoupX (72). The background soup refers to ambient RNA molecules contaminating cell-containing droplets, a common problem in droplet-based single cell RNA-sequencing technologies. Decontaminated UMIs were then filtered to discard any doublet using DoubletFinder (73). Nuclei containing more than 50% of mitochondrial UMIs were discarded. This unusually high allowance stems from the finding that cell death is part of ATC physiopathology and should not be discarded. The resulting library was scaled and normalized using the SCTransform function from Seurat. Mitochondrial content was used as a variable to regress out with SCTransform. Principal component analysis (PCA) was computed using the 3000 first variable features, and the top 30 principal components were used for SNN graph construction, clustering (resolution 1) and UMAP embedding using Seurat’s functions and Seurat recommended methods (74). Correlation analysis between genes and gene sets were computed with the fgsea function from the package fgsea v1.20.0 (75), using Spearman’s correlation coefficients between the gene of interest and each variable gene as statistics. Correlation analysis between the main axis of variation (the 3 first principal components) and genesets were computed similarly using feature loadings instead of correlation coefficients.

Cluster annotation was based on the clusters obtained from Seurat’s FindClusters function with the resolution parameter equal to 1. For all those clusters, marker genes were obtained using Seurat’s FindAllMarkers function. Clusters were annotated based on those marker genes and literature survey. Annotation was performed in two steps. In the first step only gross cell types were identified and clusters corresponding to the same gross cell type were assigned the same label. This resulted in the low resolution annotation. In the second step, each gross cell type was separated from the rest with the subset function and reprocessed as described above. The respective clusters of those reprocessed subsets were annotated as described above. This resulted in the medium and high resolution annotations. The high resolution includes all cell types, subtypes and states, while the medium resolution only includes clearly identified and relevant cell types, subtypes and states.

### Copy number alteration inference

Copy number alterations were inferred from single nuclei RNA sequencing data using InferCNV v1.10.0 (29) with default parameters and with immune and fibroblast cell types as reference. Denoised heatmaps produced by the run function were used for result interpretation and the add_to_seurat function was used to plot the HMM results on the UMAP.

### Single nuclei demultiplexing

spRNA-seq BAM files where there is no risk of sample cross-contamination were processed using picard tools v2.26.4 (76) and samtools v1.10 (77) and used to infer sample-specific single nucleotide polymorphisms (SNPs) using bcftools (77). snRNA-seq BAM files were processed using picard tools v2.26.4 and samtools v1.10, genotyped using cellSNP v0.3.2 (78) with sample-specific SNPs inferred from the matching spRNA-seq experiments, and demultiplexed using Vireo v0.5.6 (79). Vireo result tables were read in R4.1.0 and incorporated as Seurat metadata.

### TCGA data deconvolution

RNA expression profiles for the project TCGA-THCA were downloaded using the gdc-client v1.6.1 (80). Clinical variables were extracted from the supplemental materials of the THCA publication (11) and downloaded using the GDCquery command from the R TCGAbiolinks package v2.22.1 (81). Both RNA expression profiles from the TCGA and our snRNA-seq expression data annotated with cell types and filtered to match the tumor types present in the TCGA data were used as input for CIBERSORTx (82). Heatmaps were generated using the adjusted fraction matrices from CIBERSORTx with the R ComplexHeatmap package v2.10.0 (83).

### Gene regulatory network analysis

Single-cell gene regulatory network analysis was performed using pySCENIC v0.12.1 (30) with motifs v9, and database v10 with default parameters. Raw UMI counts from the whole expression matrix were used as input. The average AUC values were obtained for each medium resolution cluster by computing the mean of the individual AUC values of each cell in those medium resolution clusters.

### Mouse ESC-derived thyroid organoids experiments

Cell culture and mESC_Braf^V637E^ line generation, thyroid differentiation, Braf^V637E^ induction and the BrdU proliferation assay were performed as previously described (34). Differentiated organoids (day 25) were treated for 7 days with either differentiation medium, differentiation medium with tamoxifen for Braf^V637E^ induction, or differentiation medium with tamoxifen and 10µg/mL of soluble fibronectin protein (R&D systems, 1918-FN). Images were analyzed using QuPath v0.4.0 (65).

### Statistics

Statistical analyses and graphic representations were produced using R 4.1.0. Functions from the R base package were used unless otherwise specified and graphs were generated with R base packages or ggplot2 v3.4.2 (84).

### Study approval

This study was approved by the local ethics committee (protocol #1978).

### Data availability

The sequencing data for this study were deposited on the European Genome-Phenome Archive (EGA) under accession number EGAS00001007574. Spatial transcriptomics images at full resolution for each tumor slice are available from Zenodo with DOI <u>10.5281/zenodo.8431390</u>. R objects for single nuclei datasets and spatial transcriptomics experiments are available from Zenodo under DOI <u>10.5281/zenodo.8421521</u>. The R code used for analysis and to produce the figures of this article is available on github at https://github.com/IRIBHM-computational-groups/atc-ptc-stratification.

## Author contributions

A.T. performed experiments and computational analyses; J.R.V. performed computational analyses; M.S. performed experiments; L.C. and D.L. provided samples and pathology expertise; A.L and F.L. handled sequencing; C.M. provided expertise in thyroid cancer. M.T. provided guidance on computational analysis; S.C. and M.R. provided organoid scRNA-seq data and guidance for organoid experiments. V.D. performed computational analyses, conceived the study and collected funding. A.T. and V.D. wrote the manuscript.

## Acknowledgments

This paper is dedicated to the memory of Jacques E. Dumont. The results shown here are in part based upon data generated by the TCGA Research Network: https://www.cancer.gov/tcga. A.T. was funded by FNRS FRIA grant 1.E.019.21F, JRV by FNR Luxembourg AFR PhD grant 11587122 and Fonds David et Alice Van Buuren, Fondation Jaumotte-Demoulin and Fondation Héger-Masson, M.S. by FNRS postdoctoral grant. This research was supported by the Fondation Belge contre le cancer fundamental research grant FCC 2020-072 (V.D.), FNRS crédit de recherche J.0068.22 (V.D.) and an ULB advanced ARC grant (V.D. and S.C.).

