## supplements for "Idiosyncratic and generic single nuclei and spatial transcriptional patterns in papillary and anaplastic thyroid cancers"

### Supplemental material

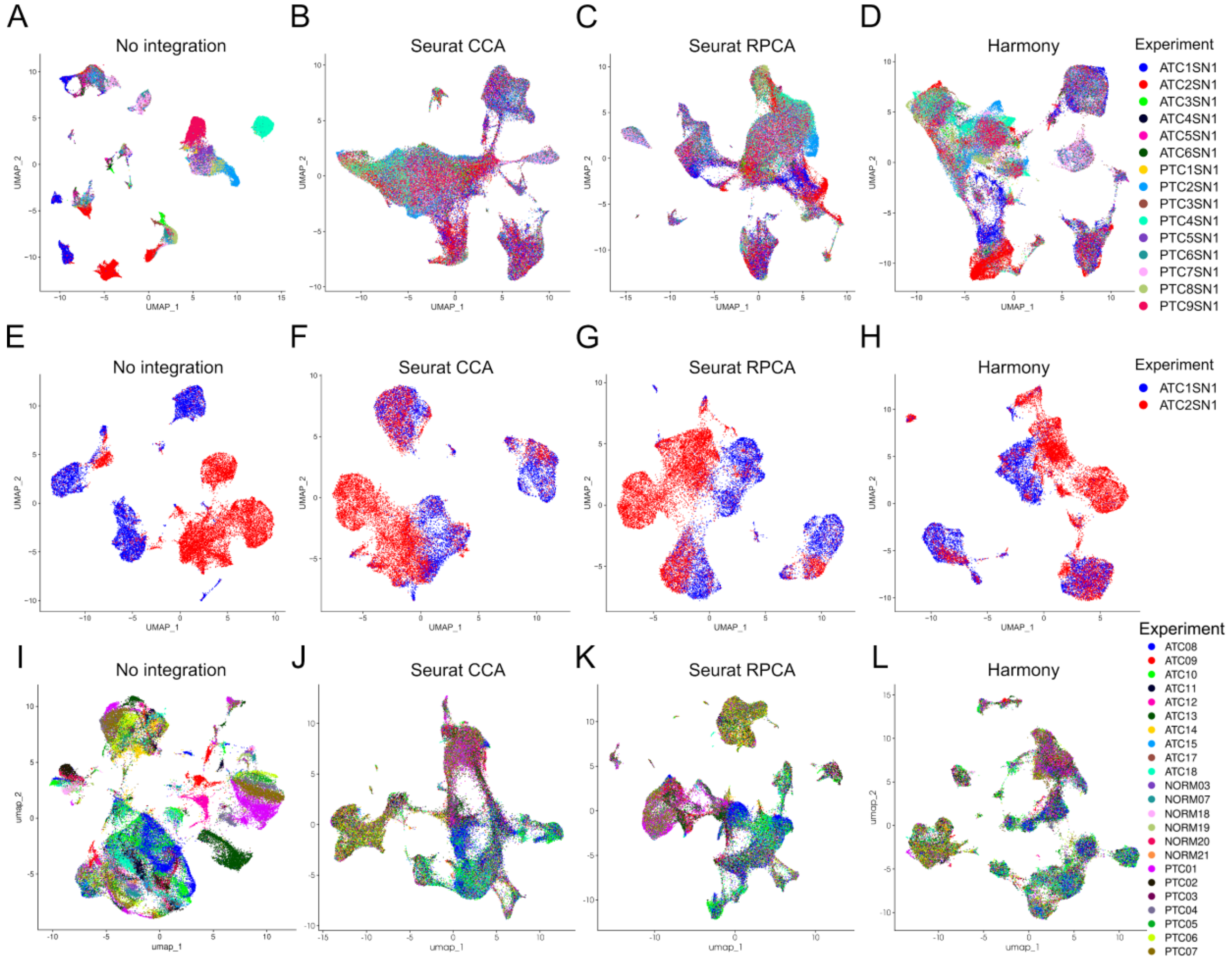


**Supplemental Figure 1. Three integration methods conceal inter-sample variation in three datasets. (A)** Umap of 9 PTC and 6 ATC samples without integration, **(B)** with Seurat's CCA integration, **(C)** with Seurat's RPCA integration, **(D)** with Harmony integration. **(E)** Umap of samples ATC1 and ATC2 without integration, **(F)** with Seurat's CCA integration, **(G)** with Seurat's RPCA integration, **(H)** with Harmony integration. **(I)** Umap of samples from Lu et al. with 6 normal, 7 PTC and 10 ATC samples without integration, **(J)** with Seurat's CCA integration, **(K)** with Seurat's RPCA integration, **(L)** with Harmony integration.

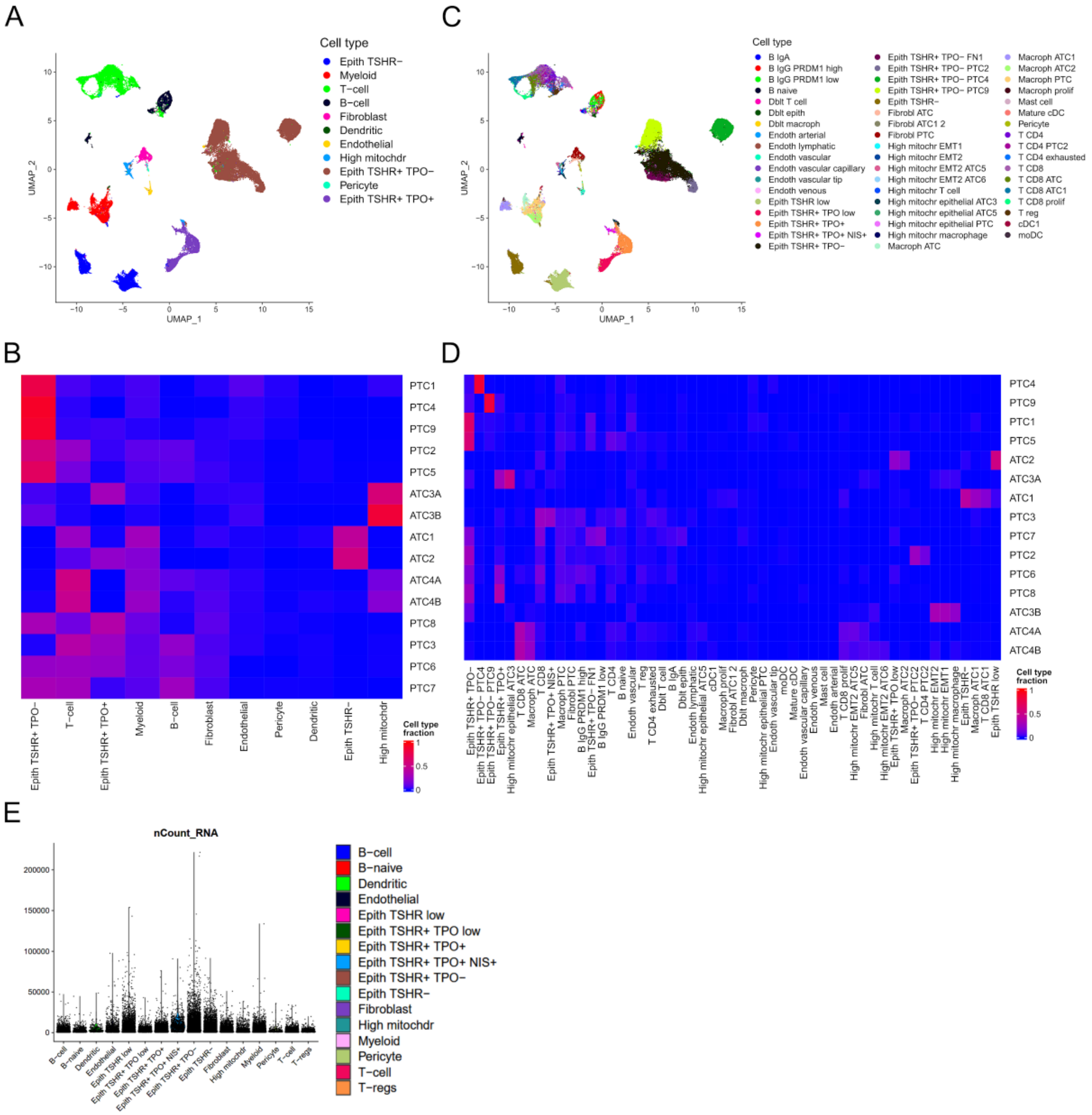

**Supplemental Figure 2. Coarse and fine grained cell types and cell states compositions. (A)** Low resolution annotation of 9 PTC and 6 ATC samples. **(B)** Low resolution annotation composition of 9 PTC and 6 ATC samples. **(C)** High resolution annotation of 9 PTC and 6 ATC samples. **(D)** High resolution annotation composition of 9 PTC and 6 ATC samples. **(E)** Number of unique molecular identifiers across cell types for 9 PTC and 6 ATC samples. Sequencing depth alone does not explain the loss of thyroid markers.

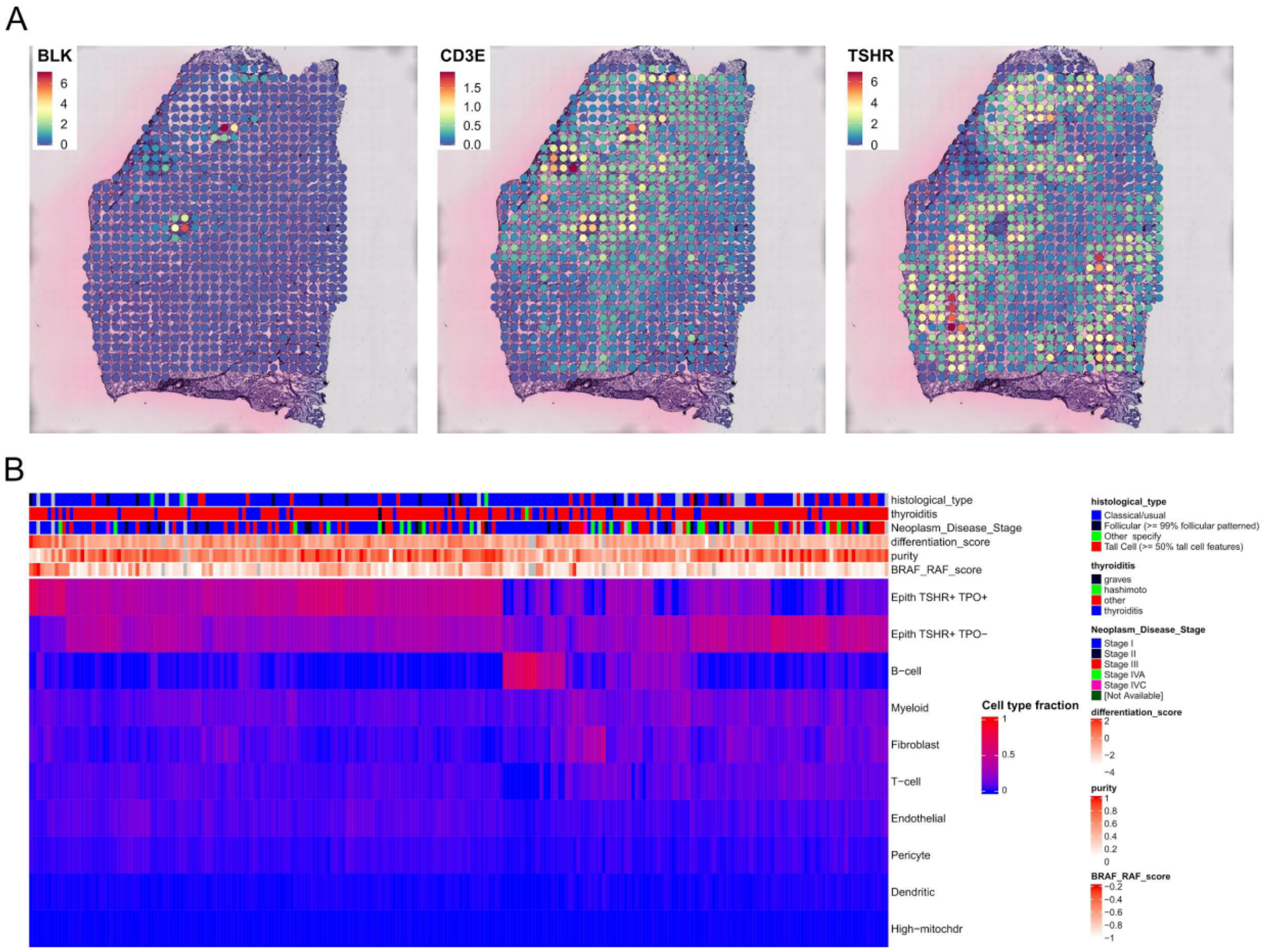


**Supplemental Figure 3. Immune cell abundance in *BRAF*$^{V600E}$ PTC. (A)** Expression level of immune and epithelial markers in PTC6S1. **(B)** Cibersortx-inferred cell type composition of 234 *BRAF*$^{V600E}$ mutated TCGA samples.

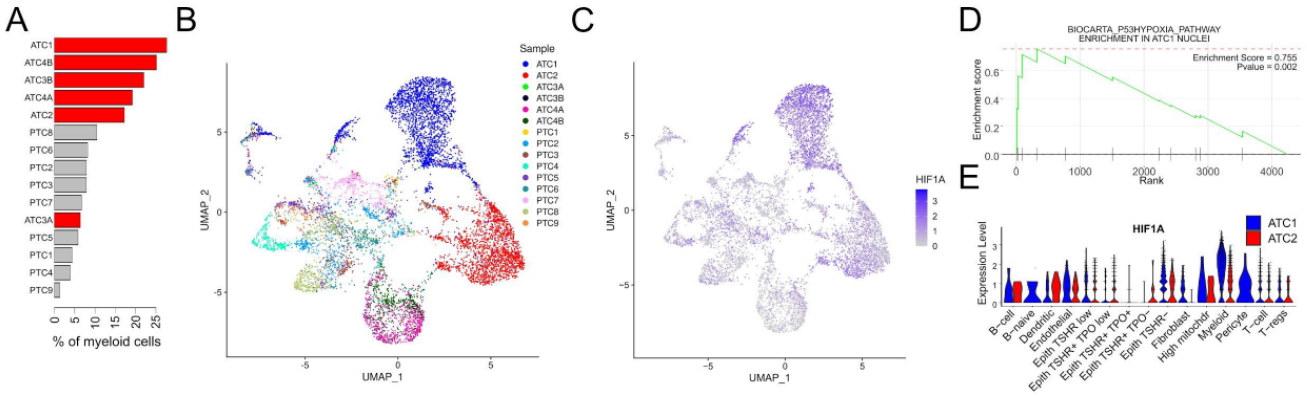


**Supplemental Figure 4. An idiosyncratic myeloid cell state is associated with hypoxia in an ATC. (A)** Fraction of myeloid cells in snRNA-seq samples. Red depicts ATCs. **(B)** UMAP of myeloid cells with sample annotation. Cells from ATC1 and ATC2 form distinct clusters. **(C)** Expression of *HIF1A* gene in myeloid cells. It is increased in ATC1. **(D)** Enrichment for hypoxia pathway in ATC1 myeloid nuclei. **(E)** Expression level of gene *HIF1A* across cell types for samples ATC1 and ATC2. *HIF1A* is increased in all cell types present in ATC1, pointing to systemic hypoxia.

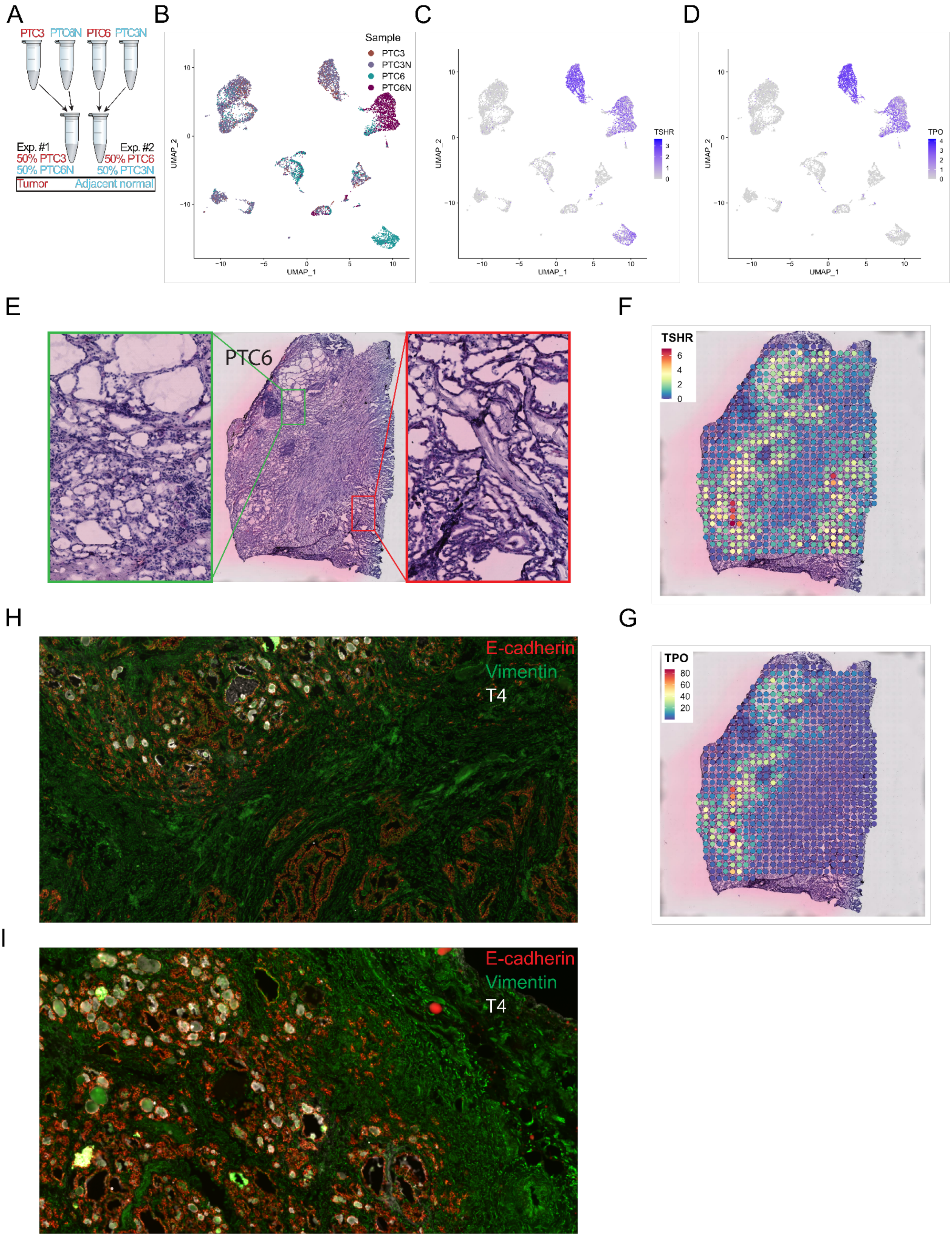


**Supplemental Figure 5. A cell mixing experiment resolves the status of TPO$^+$ cells.** Tumor slices could contain adjacent normal nuclei, raising doubts about the status of TPO$^+$ cells: are they surprisingly well differentiated cancer cells or normal contaminants? As our original study design did not explicitly include normal samples, we performed additional experiments mixing tumor and adjacent healthy tissues to settle this question. **(A)** Schematic representation of the nuclei mixing experiment for samples PTC3, PTC6 and their adjacent normal tissue. **(B)** UMAP showing the sample of origin of each nuclei as assigned by Vireo for samples PTC3, PTC6 and their adjacent normal tissue. **(C, D)** Expression level of *TSHR* (**C**) and *TPO* (**D**) in PTC3, PTC6 and their adjacent normal tissue. Epithelial nuclei from PTC3, PTC3N and PTC6N express *TPO* and *TSHR*, while those from PTC6 only express *TSHR*. This is corroborated by spatial data. **(E)** H&E image of PTC6 with zoom on the hormone-producing area (left) and the PTC-like area (right). **(F, G)** Expression level of *TSHR* (**F**) and *TPO* (**G**) in PTC6.**(H)** Immunofluorescence staining of PTC6 at the intersection between hormone-producing and PTC-like areas. T4 staining (white) is consistent with the *TPO* expression pattern of panel **G**. **(I)** Immunofluorescence staining of PTC6 normal adjacent sample.

Ninety six percent of $NIS^+$ nuclei (a subset of $TPO^+$ nuclei) belonged to PTC3 and 23% of $TPO^+$ nuclei to PTC6 (compare Figure 1A and Figure 2A). Of note, all epithelial nuclei from the PTC3 sample were $NIS^+$, while PTC6 included both $TPO^+$ (19% of PTC6) and $TPO^-$ (17% of PTC6) nuclei. The cancer status of the latter is demonstrated by their membership to the main PTC cancer cell cluster (compare Figure 1A and Figure 2A). Even minimal batch effects could confound the comparison of cells with subtle differentiation differences. Thus, to ascertain the tumoral/normal status of our $NIS^+$ and $TPO^+$ nuclei while controlling for potential batch effects, we devised two cell mixing experiments with both patients' tumor and adjacent normal samples (Supplemental Figure 5A). The first experiment mixed nuclei from PTC3 and from PTC6N, a non-cancer block adjacent to PTC6. Conversely, the second experiment mixed nuclei from PTC6 and from PTC3N. As previously, nuclei from the two experiments were pooled in silico and processed as a single dataset, without batch effect correction, and patient identities were recovered with Vireo. Based on the sample of origin of each nuclei (Supplemental Figure 5B), *TSHR* (Supplemental Figure 5C) and *TPO* (Supplemental Figure 5D) expression, we observed that epithelial nuclei were subdivided in 3 clusters. All epithelial nuclei from PTC3 and PTC3N were homogeneously mixed in the same most differentiated epithelial cluster (Supplemental Figure 5D). Conversely, epithelial nuclei from PTC6 (tumor sample) and PTC6N (normal sample) were subdivided in 2 clusters : one less differentiated and exclusively from the tumor sample and one more differentiated from both normal and tumor samples. This was consistent with the presence of a mix of normal and malignant nuclei in the tumor sample of PTC6. But it also indicated that PTC3–to our surprise—did not contain malignant nuclei despite its highly disturbed morphology. The normal/malignant dichotomy in PTC6 can also be observed in the spRNA-seq experiment where normal nuclei ($TPO^+$) are separated from the tumoral nuclei ($TPO^-$) by a fibrous area (Supplemental Figure 5E-H). Immunofluorescence also confirmed that the $TPO^+$, follicular shaped area produces T4, while the $TPO^-$ papillary shaped area does not (Supplemental Figure 5H), and that PTC6N is only composed of follicular, T4-producing structures (Supplemental Figure 5I). These results ascertain the non-cancer status of $TPO^+$ nuclei. Some nuclei expressing *TPO* at lower levels were also present in ATC2. ATC2 spRNA-seq slices do include residual follicular structures expressing *TPO*.

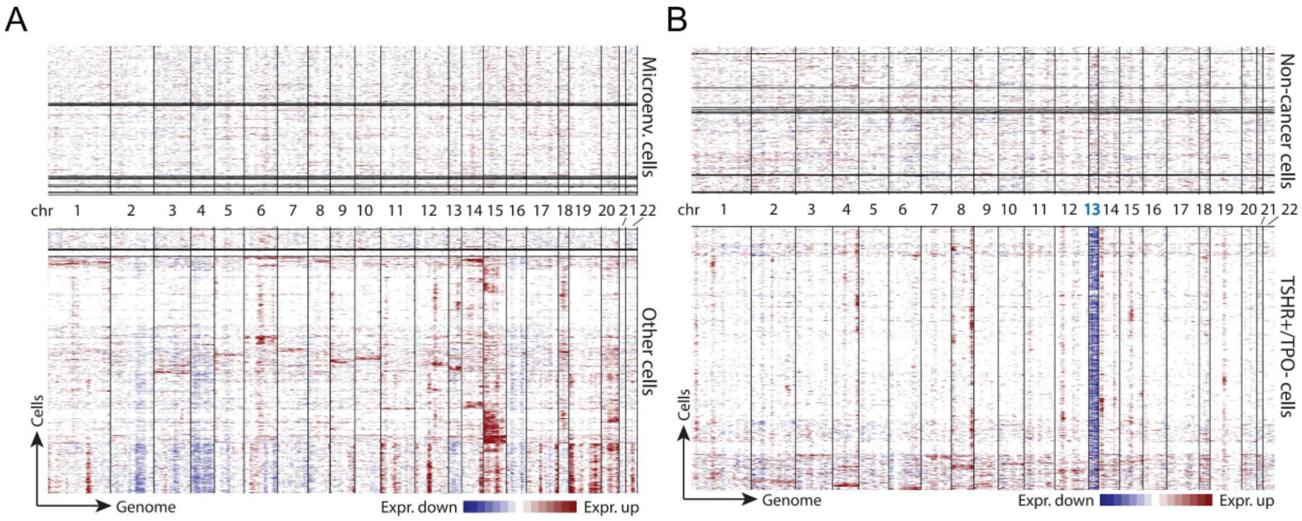


**Supplemental Figure 6. DNA copy number profiles of *TSHR*$^{low/-}$ cells. (A)** DNA copy number alterations inferred from snRNA-seq by infercnv in sample ATC1. **(B)** DNA copy number alterations inferred from snRNA-seq by infercnv in sample PTC4.

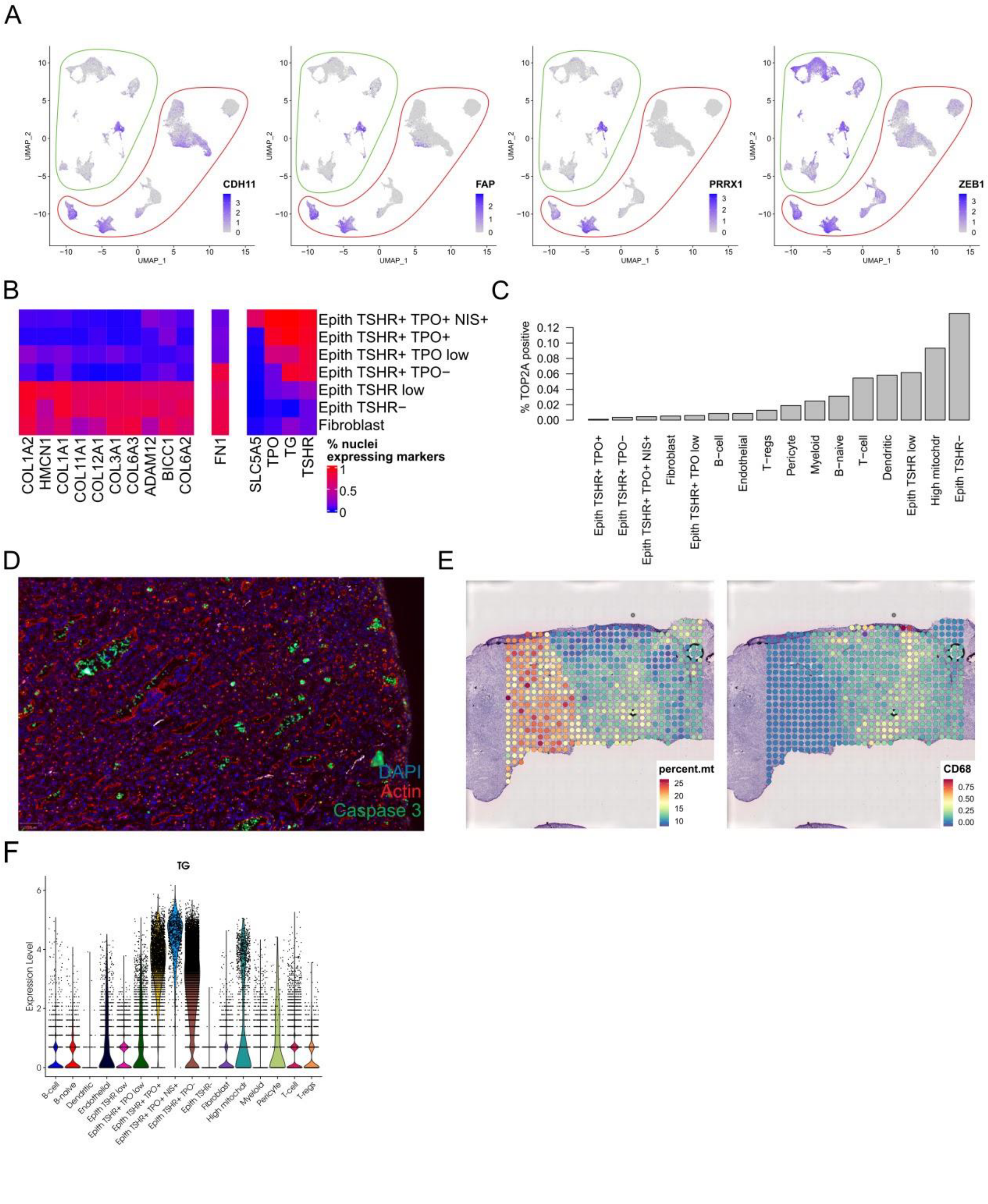


**Supplemental Figure 7.** ***TSHR*$^{low/-}$ cells, EMT and cell death. (A)** Expression level of mesenchymal markers in 9 PTC and 6 ATC samples. The microenvironment is highlighted in green and nuclei of epithelial origin in red. **(B)** Fraction of cells expressing mesenchymal and thyroid differentiation markers in the fibroblast and epithelial subsets. **(C)** Fraction of *TOP2A* positive nuclei per cell type. **(D)** Immunofluorescence staining of ATC3 showing cells positive for the activated form of Caspase3. Apoptotic cells stained for Caspase3 are primarily located in the follicular lumens. **(E)** Expression level of mitochondrial genes and *CD68* in sample ATC3BS1. **(F)** Expression level of *TG* for all subsets of the medium resolution annotation in 9 PTC and 6 ATC samples. It demonstrates that apparent TG expression in fibroblasts panel B is likely a non cell type-specific effect of ambient TG RNA contamination.

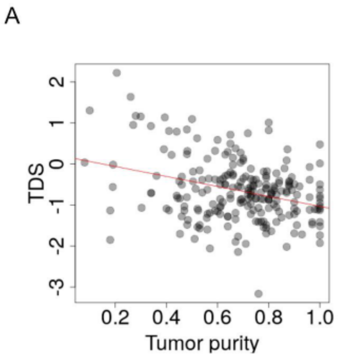


**Supplemental Figure 8. Correlation between TDS and purity in the TCGA cohort. (A)** Scatterplot of the thyroid differentiation score (TDS) vs. tumor purity in the BRAF-mutated PTC samples of the TCGA cohort. Spearman's correlation between TDS and purity is $\rho = 0.32$, $p < 2x10^{-16}$.

**Supplemental tables 1 to 9 are available online.**

**Supplemental table 1. Sample summary.**

**Supplemental table 2. snRNA-seq quality metrics summary.**

**Supplemental table 3. spRNA-seq quality metrics summary.**

**Supplemental table 4. Cell type markers at low resolution.**

**Supplemental table 5. Cell type markers at medium resolution.**

**Supplemental table 6. Cell type markers at high resolution.**

**Supplemental table 7. Composition of previously published PTC cohorts.**

**Supplemental table 8. References of antibodies used for immunostaining.**

**Supplemental table 9. SCENIC AUC values averaged by medium resolution clusters.**
